# Common and distinct BOLD correlates of Simon and flanker conflicts which can(not) be reduced to time-on-task effects

**DOI:** 10.1101/2023.05.19.541457

**Authors:** Jakub Wojciechowski, Katarzyna Jurewicz, Patrycja Dzianok, Ingrida Antonova, Katarzyna Paluch, Tomasz Wolak, Ewa Kublik

## Abstract

The ability to identify and resolve conflicts between standard, well trained behaviors, and behaviors required by the current context is an essential feature of cognitive control. To date, no consensus has been reached on the brain mechanisms involved in exerting such control: while some studies identified diverse patterns of activity across different conflicts, other studies reported common resources across conflict tasks or even across simple tasks devoid of conflict component. The latter reports attributed the entire activity observed in the presence of conflict to longer time spent on the task (i.e. to the so-called time-on-task effects). Here we used an extended Multi-Source Interference Task (MSIT) which combines Simon and flanker types of interference to determine shared and conflict-specific mechanisms of conflict resolution in fMRI, and their separability from the time-on-task effects. Large portions of the activity in the dorsal attention network and decreases of activity in the default mode network were shared across the tasks and scaled in parallel with increasing reaction times. Importantly, activity in the sensory and sensorimotor cortices, as well as in the posterior medial frontal cortex (pMFC)–a key region implicated in conflict processing–could not be exhaustively explained by the time-on-task effects.

## 1. Introduction

A large part of our everyday behavior is guided by well-learned reactions and pre-existing associations between stimuli and responses. However, a truly effective behavior depends on the ability to identify and resolve conflicts between our standard reactions and those required by the current context. The mechanisms of cognitive control that enable such flexible behavior have frequently been studied with response conflict tasks, i.e. tasks which use various input stimuli to simultaneously prompt different, competing responses. A popular example of a paradigm which elicits response conflict is the so-called Simon task (Simon 1967), in which conflict occurs due to the spatial arrangement of stimuli and responses. In the most classic form of the task, subjects respond to a feature of a stimulus (e.g., color) presented to either the left or right of a central fixation, by pushing response buttons with their left (e.g., for blue) or right (e.g., for red) hand. Performance is impaired when the side of presentation does not match the side of response, (e.g., a red stimulus presented on the left side). Another frequently used paradigm is the flanker task (Eriksen and Eriksen 1974), in which conflict occurs due to a spatial cluttering of symbols. In its classic form, subjects have to respond to the central letter, number or arrow, while ignoring the surrounding distractors. Most pronounced interference is observed when distractors are associated with alternative responses. These two tasks represent the examples of the stimulus-response (Simon) and stimulus-stimulus (flanker) interference. While both eventually evoke the response conflict, i.e., increased competition between behavioral alternatives, the origin of interference is different in the two types of task. It stems from a direct mismatch between spatial properties of the stimuli and response dimension in Simon; and begins already at the level of stimulus interpretation in the flanker task (Egner et al. 2007).

Over the past few decades, research on cognitive control has tried to determine if different conflicts are resolved through separate mechanisms that operate in a conflict-specific manner or if there is a central resource for dealing with all kinds of conflicts. However, previous imaging studies which compared conflict-induced activity across different tasks brought rather divergent results. Several studies reported distinct regions involved in resolving different forms of cognitive interference (Egner et al. 2007; van Veen and Carter 2005). Others found substantial differences and some overlap of activation between conflicts (Fan et al. 2003; Frühholz et al. 2011; Liu et al. 2004; Wager et al. 2005; Wittfoth et al. 2006). There were also studies which found mostly overlapping mechanisms underlying resolution of different conflicts (Peterson et al. 2002). Multiple factors could possibly account for such heterogeneity of results. Compared tasks were often based on different types of stimuli, making it more difficult to disentangle conflict-related effects from task-specific processing of information. Also, the early studies were often severely underpowered which might have led to either a decreased sensitivity for task-related activations or an increased number of false positive findings (as demonstrated in Button et al. 2013; Ioannidis 2005; Eklund et al. 2018; Poldrack et al. 2017; Szucs and Ioannidis 2020). The meta-analytic approach indicated that some limited regions, including the posterior medial frontal cortex (pMFC), activate consistently across different conflict tasks, while some consistent between-task differences were also identified in fronto-parietal regions (Cieslik et al. 2015; Li et al. 2017; Nee et al. 2007). Recently, however, the conflict-processing role of regions commonly engaged by the conflict tasks has been questioned from another perspective. A new important stand on the correlates of conflict processing, as observed in fMRI research, came from the series of studies which investigated reaction time (RT)-related variability of the blood-oxygen-level-dependent (BOLD) signal (Carp et al. 2010; Grinband et al. 2008; Grinband et al. 2011; Weissman and Carp 2013; Yarkoni et al. 2009). It has been shown that activity in a large set of regions, including pMFC, changes as a function of RT across various experimental designs (Grinband et al. 2008; Yarkoni et al. 2009). In particular, even the most simple task which does not require any selection (i.e. a simple button press in response to stimulus appearance) demonstrates close correlation between the RT and the strength of the blood-oxygenation-level-dependent (BOLD) response (Weissman and Carp 2013). In response-conflict tasks, conflict conditions are always characterized by longer RT in comparison to control, which does not entail interference. Thus, RT constitutes a confounding factor, which may account for a large part of brain activity observed in these tasks. While it is true that RTs are longer *because of* the occurrence of conflict, brain activity associated with longer RTs may not necessarily reflect conflict-related processing. For example, a slower response might be accompanied by a longer exposure to visual stimuli, prolonged need for attention maintenance, increased need for task rehearsal etc., all of which may contribute to the changes in the BOLD signal in different parts of the brain. The question posed by the recent neuroimaging research was whether conflict-related activity can be distinguished from the effects of the time-on-task as such.

To address this question Carp et al. 2010 used a Multi-Source Interference Task (MSIT) which includes both Simon- and flanker-type interference, combined in a single condition to maximize the strength of conflict and recruitment of the related brain resources (Bush et al. 2003). The study showed that RT-related variability of the signal in no-conflict trials of this task sufficed to predict the entire activity observed in multi-source interference trials when BOLD responses were appropriately scaled on the basis of RT. Another study showed that even a paradigm with only one possible response (i.e. a simple button press at stimulus appearance that does not require any perceptual or motor selection) can be successfully used to model multi-source conflict activity in MSIT with the same scaling method (Weissman and Carp 2013). Similarly, there was no difference in BOLD responses to MSIT conflict and no-conflict conditions when trials from each category were subsampled to have similar RT (Carp et al. 2010). The same results were obtained also with the Stroop task (Grinband et al. 2011). As such, these studies strongly suggest that the resources recruited by conflict tasks are shared across cognitive operations which influence RT, limiting their interpretation in terms of conflict processing. The so-called time-on-task effects appeared more closely tied to task maintenance or attention that may play a role in even the simplest tasks.

The exhaustive role of RT-related effects in interference tasks is hard to reconcile with the large number of reports of pMFC engagement in various aspects of conflict processing (Ullsperger et al. 2014). It also remains at odds with findings based on electrophysiological techniques, which show conflict-related encoding in the same regions that seem to reflect pure time-on-task effects in fMRI data (Sheth et al. 2012; Ebitz et al. 2020). However, the all-encompassing time-on-task effects are also perplexing on the grounds of fMRI research. The time-on-task effects occur independently from the presence of conflict (i.e., across all task conditions). Therefore, while regions commonly involved in response conflict may or may not reflect the time-on-task effects, conflict-specific activity would need to diverge from the RT-based predictions. If different types of conflicts have different sources of interference, RT-related variability should not explain the entire signal that is related to processing of conflict.

In our study, we wanted to reiterate the question about common and conflict-specific mechanisms of conflict processing. We used MSIT, the paradigm previously adopted for demonstrating the exhaustive role of time-on-task effects in BOLD responses within conflict tasks, but in the extended version comprising also single conflict conditions. The use of a single type of stimuli and the same instruction for probing Simon and flanker conflicts, in a task performed by the same participants in a single session, minimized the role of task-specific attributes. Importantly, common and conflict-specific responses for the conditions of extended MSIT have recently been identified with magnetoencephalography (MEG) (Wiesman and Wilson 2020; Wiesman et al. 2020), giving a hint that there are at least partially separate mechanisms underlying resolution of these two conflicts. Thus, we used a task that is ideally suited for contrasting different types of conflicts, as well as for determining the contribution of time-on-task effects to clarify some of the longstanding questions on the common and distinct mechanisms involved in the response-conflict resolution.

## 2. Materials and methods

### 2.1 Participants

Forty-one (22 female, 19 male) healthy young adults (mean age = 24.6 years, SD = 4.2) participated in the study. One subject was excluded due to incomplete data. All subjects were right-handed and had normal or corrected-to-normal vision. Subjects were recruited at the University of Warsaw and via social media. Before taking part in the study, all subjects gave their written informed consent for participation and were screened for any contraindications to magnetic resonance imaging. The study was approved by the ethical committee of the University of Nicolaus Copernicus in Toruń, Poland, and was conducted in accordance with the Declaration of Helsinki.

### 2.2 Experimental procedure

The task used in the study was an extended version of the Multi-Source Interference Task (MSIT; Bush et al. 2003). Originally, MSIT consisted of two conditions: no-conflict and multi-source conflict trials. The latter combined effects of stimulus-response incongruence (Simon effect) and stimulus-stimulus incongruence (flanker effect). In the extended version which we use here, these two types of cognitive interference were also included separately, with only one source of interference (Simon or flanker) present at a given trial.

The three digits (from the set 0, 1, 2 and 3) were presented in the center of the screen: two of them were the same and one (digit 1, 2 or 3) was always different (Figure 1B). The stimuli were presented on the dark screen (RGB: 48, 48, 48), the color of the stimulus was light gray (RGB: 226, 226, 226). Subjects had to identify the unique digit (target) and indicate it by responding with the appropriate finger of the right hand, i.e., with index finger for “1”, middle finger for “2”, or ring finger for “3”. In the no-conflict trials (00) the position of the unique digit was congruent with the correct button choice and the target was flanked by zeros, which were not mapped to any response (e.g., “003”). In Simon-only condition (S0) the target was flanked by zeros, but the position of the unique digit was incongruent with the spatial position of the correct response (e.g., “030”). Conversely, in flanker-only condition (F0), the unique digit and correct response were spatially congruent, but the flankers were always different from zero, i.e., entailed their own possible button-choices (e.g., “223”). In the multi-source interference condition (FS) both the position of the unique digit and response-mapping of the flankers were incongruent with the correct response associated with the target (e.g., “232”).

**Figure 1:**
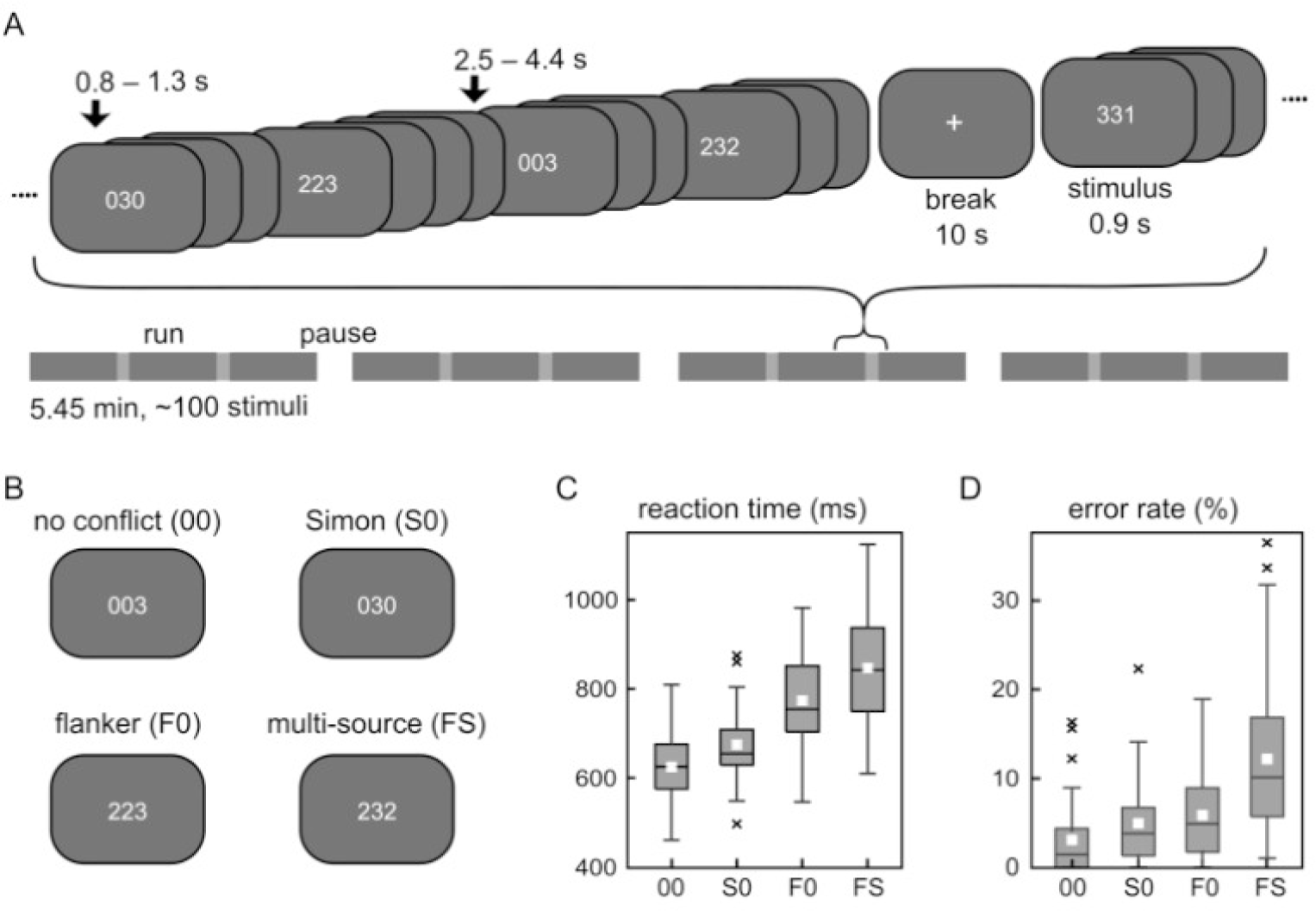
MSIT experimental paradigm. A) Stimuli were presented for 900 ms with a standard interval of 800-1300 ms after a stimulus and a longer interval of 2500-4400 ms after a sequence of 3 or 4 stimuli of the same category (miniblock), with two additional 10 ms intervals per run. The whole session included ∼400 stimuli and was divided into four runs. B) Example stimuli in no-conflict (00), Simon (S0), flanker (F0) and multi-source (FS) conflict conditions. In each case of the example, “3” is the correct response and requires a button press with a third (ring) finger. C) Reaction time (for correct responses) and D) error rate in the four MSIT conditions. White rectangles represent the mean, boxes span the 25th to 75th percentile of the distribution with the median marked as a horizontal bar, crosses represent the outliers.

Trials were presented for 900 ms in event-related design with additional grouping into mini-blocks of three or four consecutive trials (equal number) drawn from the same condition, to further increase the power of contrasts between conditions. Each trial was followed by 800-1300 ms inter-trial interval within and 2500-4400 ms between mini-blocks. Three 10-second pauses presenting a fixation cross were inserted after each third of a run. Order of stimuli was pseudo-random, such that there were no more than two consecutive mini-blocks of the same condition type. On average, a single fMRI run of the task consisted of approx. 100 trials (∼25 per condition). Subjects underwent four acquisition runs, each lasting 5:45 minutes (see Figure 1A for the diagram of an experimental timeline).

The experimental protocol (stimulus delivery and response registration) was controlled by the Presentation software (Neurobehavioral Systems Inc.). Stimuli were presented on MR-compatible full HD display screen (NordicNeurolab Inc.) set behind the MR scanner. A mirror system mounted on a head coil was used to project the visual stimuli to the subject. Behavioral responses were collected using MR-compatible response pads (SMIT-Lab Co.). Before entering MR and beginning the experiment, subjects underwent brief training (lasting on average 2:20 minutes) of the task to familiarize them with the procedure.

### 2.3 Behavioral data analysis

From reaction time and fMRI analyses we discarded trials with incorrect, preemptive (< 200 ms, n = 2 across all trials and subjects) or very slow responses (> 1700 ms, exceeding shortest inter-stimulus interval, n = 19 across all trials and subjects). Mean RT for correct trials and error rate for 00, S0, F0 and FS trials for each subject were entered in 1×4 repeated-measures model ANOVA. To check if Simon and flanker have additive effects on behavior or interact with each other when present simultaneously in multi-source conflict condition, we tested whether RT and accuracy changes are equal in FS versus summed F0 and S0 conditions (i.e., whether FS - 00 = (F0 - 00) + (S0 - 00)). Pairwise comparisons applied Tukey’s HSD (honestly significant difference) test and held the familywise error rate at 0.05. All statistical analyses were performed in the MATLAB environment and JASP software (JASP Team, 2023).

### 2.4 Image acquisition

Scanning was performed on a 3T Siemens Prisma MRI scanner (Siemens Medical Solutions Inc.) using a 64-channel RF head coil. Structural MRI data were acquired with a T1-weighted 3D MP-Rage sequence (TR = 2.4 s TI = 1000 ms, TE = 2.74 ms, 8° flip angle, matrix 320×320, FOV = 256×256 mm, isometric voxel size 0.8 mm, 240 slices), lasting 6:52 minutes. Functional data were acquired using Multi-band (Siemens naming: Simultaneous Multi-Slice) EPI sequence (TR = 750 ms, TE = 31ms, FA = 52°, FOV = 220×220 mm, matrix = 110×110, 72 axial slices, isometric voxel size 2 mm, no interslice gap, Slice Accel. Factor = 8, In-Plane Accel. Factor = 1, IPAT = 8).

### 2.5 Image preprocessing

Preprocessing of the MRI data was performed using SPM12 (SPM; WellcomeTrust Centre for Neuroimaging, London, UK), FSL (Smith et al. 2004) and in-house MATLAB code. Multi-band EPI sequence provides an overall signal-to-noise increase of 40%, but introduces image distortions due to the accumulation of phase errors in the k-space. To account for this effect, all functional scans had an even number of runs with the opposite phase coding direction (Anterior-Posterior and Posterior-Anterior). Spatial distortions were corrected with the FSL *topup()* tool based on the average representation of the images in both directions. Functional data was spatially realigned to the mean image, to which the structural image was then co-registered. Segmentation and normalization to the common MNI space was performed based on high-resolution structural images with resampling to 1 mm isometric voxels. The obtained transformation parameters were applied to the functional volumes with resampling to 2 mm isometric voxels. The normalized functional images were spatially smoothed with a Gaussian kernel of full-width half-maximum (FWHM) of 5 mm, and additionally high-pass filtered above 0.008 Hz (time constant of 128 sec).

### 2.6 Image analysis

#### 2.6.1 Model estimation

Functional images were analyzed using the general linear model in SPM12. BOLD responses evoked by each of the four types of trials included in MSIT (00, S0, F0, FS) were modeled as separate regressors along with their parametric modulators, which reflect a trial-by-trial RT-BOLD relationship within the given condition. Each RT-modulator was orthogonalized in relation to the condition regressor, as implemented in SPM (in our design this is equivalent to demeaning of the RTs; Mumford et al. 2015). Incorrect trials were modeled as a separate regressor and discarded from subsequent analyses. The model also included six motion-related nuisance regressors and one regressor for pauses. Whole-brain group-level analyses used a voxel-level threshold of p < 0.001 for each side of comparison (as per standard SPM analysis) and a cluster level threshold of pFWE = 0.05, unless stated otherwise.

#### 2.6.2 Assessment of activity induced by conflicts

To demonstrate which regions were engaged by different types of interference, we first compared each of the conflict conditions to a no-conflict baseline (i.e., two sided contrasts were defined: [S0 vs 00], [F0 vs 00] and [FS vs 00]). We treated the three conflict conditions separately (and not within a full factorial design) for several reasons: (1) one of the main objectives of this study was to test how RT-related variability explains conflict signals; RT is monotonically increasing across task conditions, enabling RT-based predictions for each type of conflict, (2) another objective of the study was to test conflict-specific effects distinguishing Simon and flanker conflict processing; any interaction between Simon and flanker effects would impact the estimates (main effects) of Simon- or flanker-related activity, (3) previous imaging studies investigating the time-on-task effects in MSIT used a version of task with multi-source conflict only; testing FS vs 00 contrast enables comparisons between current and previous reports. However, for completeness, the interaction effects between the conflicts were also tested in a separate contrast, which compared the summed changes of the BOLD signal in Simon and flanker conditions with the changes observed in multi-source conflict (i.e., testing whether FS - 00 = (F0 - 00) + (S0 - 00)), which can be transformed to [(FS + 00) vs (S0 + F0)] contrast).

Additional analyses were performed to specify which of the observed effects were common across Simon and flanker conditions, and which activations resulted from only one of the conflicts. First, we performed conjunction analysis of Simon and flanker activations, [(S0 vs 00) ∩ (F0 vs 00)], to specify which regions engage both in Simon and in flanker conflicts. Conjunction was based on (S0 vs 00) and (F0 vs 00) maps, which were thresholded at lower than regular voxel-level threshold of p < 0.01 to include more regions engaged by the task, even if they differed in activation strength. The resulting conjunction map was thresholded again at a standard voxel-level threshold of p < 0.001. Second, we directly compared Simon and flanker conditions [F0 vs S0] to reveal regions of maximum difference in the activity evoked by these two conflicts. Third, to expose qualitative rather than quantitative differences, we masked out the commonly modulated regions (i.e., conjunction result, [(S0 vs 00) ∩ (F0 vs 00)]) from the map of the two-sided contrast of Simon and flanker conditions ([F0 vs S0]). This operation allowed us to obtain regions specifically more active in flanker than in Simon condition, expressed in positive contrast values, and specifically more active in Simon than in flanker condition, expressed in negative contrast values.

#### 2.6.3 Assessment of activity exceeding time-on-task effects

To test if activity observed in MSIT conditions is associated with the time-on-task or conflict-related effects, we performed RT-regression analyses proposed in previous works (Carp et al. 2010; Weissman and Carp 2013). First, we estimated the time-on-task effects in Simon, flanker, and multi-source conflicts on the basis of their mean RT and GLM model parameters obtained for no-conflict condition. Namely, we used the beta estimate of the mean activity in no-conflict condition (00) and the beta estimate of the linear change of signal with RT in no-conflict condition (parametric modulator of 00 trials, 00_RTmod_, Suppl. Figure S1A) to predict the level of activity based on mean RT change in a conflict condition (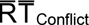) with respect to mean RT in the no-conflict condition, 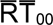). Thus, for each voxel, activity was estimated according to the following formula (separately for each conflict condition):

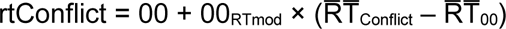

Next, for each of the conflict conditions we compared their activity maps and the corresponding RT-predicted counterparts (i.e., two-sided contrasts were performed: [S0 vs rtS0], [F0 vs rtF0] and [FS vs rtFS]). These contrasts showed regions of activity not fully explained by (i.e. exceeding) the time-on-task effects.

A recent work focused on discerning time-on-task from main task conditions effects (preprint Mumford et al., 2023) proposed to adopt a model, in which all conditions share a single, non-orthogonalized RT-modulator. Such a design would remove all differences resulting from the time-on-task from the comparison of the task conditions. However, this approach is only applicable, when there is no condition-RT interaction (i.e. condition-specific RT modulation). This is not the case in the extended MSIT, where we found a significant conflict-RT interaction (Suppl. Figures S1 and S2). The method proposed by Mumford and colleagues offers increased power, but is not applicable in the case of conflict-RT interaction, in which the approach proposed by Carp and colleagues serves the same purpose, without conflating the basic time-on-task effect and conflict-specific RT-variability.

One other approach for accounting for the time-on-task effects in the conflict analysis requires RT-matching of trials between the conditions (Beldzik et al. 2015, Carp et al. 2010). This approach is also not well suited for the extended MSIT: reaction times for the no-conflict and multi-source conflict conditions are too different to enable subsampling a sufficient number of RT-matched trials. Thus, we made a supplementary RT-matched comparison only for the two uni-source conflicts (Simon and flanker). To match the two RT distributions we exclude on average 23% of fastest S0 and slowest F0 trials for each participant.

In parallel to analyses provided for entire Simon- and flanker-related activity, we performed conjunction analysis of activity exceeding the time-on-task effects in Simon and flanker conditions, [(S0 vs rtS0) ∩ (F0 vs rtF0)]. Again, for this conjunction analysis, we lowered the significance threshold of the constituent maps to uncorrected voxel-level p < 0.01 to disregard regional differences resulting merely from the strength of the effect. We also directly compared the strength of Simon- and flanker-related activity unexplained by time-on-task, [(F0 vs rtF0) vs (S0 vs rtS0)]. Lastly, to highlight spatially distinct regions resulting from RT-regression in Simon and flanker conditions, we masked the difference of activity exceeding time-on-task in the two conflict conditions [(F0 vs rtF0) vs (S0 vs rtS0)], by their conjunction [(S0 vs rtS0) ∩ (F0 vs rtF0)].

#### 2.6.4 ROI analysis

Lastly, we performed additional analyses in the pMFC, to verify the contribution of time-on-task effects in this region, most often implied in conflict processing and selected for similar analyses in previous studies (Carp et al. 2010; Weissman and Carp 2013). Spherical regions of interest (ROI) with a 8 mm radius were centered at the peak of activity identified in pMFC in the conflict vs no-conflict conditions (S0 vs 00, F0 vs 00 and FS vs 00). For most direct comparison with previous studies, we also analyzed the observed and RT-predicted signals evoked by multi-source conflict condition in ROIs centered at the peaks identified by Carp et al. 2010 and Weissman and Carp 2013. Thus, all ROIs were selected orthogonally to the time-on-task effects in question. To outline the dependencies between the observed effect sizes and the study sample, we sub-sampled individual beta weights for the observed and RT-predicted signal (N = 24, 1000 randomly drawn samples). This simulation illustrated the chances of observing a “significant” result (p < 0.05) for the smaller study sample (as used in the previous studies).

## 3. Results

### 3.1 Behavioral indices of conflict effects in MSIT

The analysis of RTs from correct trials (Figure 1C) showed that there were significant differences between the task conditions (F(3,117) = 250.43, p < 0.001, η2 = 0.86). Relative to no-conflict trials (M = 625.13 ms, SD = 78.17), participants were significantly slower in the Simon (M = 675.52 ms, SD = 81.16, p < 0.001), flanker (M = 774.17 ms, SD = 92.17, p < 0.001), and multi-source (M = 847.96 ms, SD = 119.64, p < 0.001) conflict conditions. Furthermore, participants were significantly slower in flanker than in Simon condition (p < 0.001). The multi-source conflict also resulted in slower responses than in both Simon (p < 0.001) and flanker (p < 0.001) conditions.

Similarly, the analysis of error rate (Figure 1D) revealed significant differences between task conditions (F(3,117) = 51.60, p < 0.001, η2 = 0.57). Relative to no-conflict trials (M = 3.15%, SD = 4.21), participants had a significantly higher error rate in the Simon (M = 5.00%, SD = 4.83, p = 0.003), flanker (M = 5.91%, SD = 5.00, p < 0.001), and multi-source (M = 12.22%, SD = 8.92, p < 0.001) conflict conditions. However, accuracy was not significantly different between flanker and Simon trials (p = 0.276), while the multi-source condition resulted in higher error rate than in either of the single conflicts (p < 0.001).

Previous studies suggested that Simon and flanker conflicts may interact with each other when simultaneously present in multi-source condition (Wiesman and Wilson 2020). To check if Simon and flanker have additive or interactive effects on behavioral performance, we tested whether RT and error rate changes are equal in multi-source and the sum of Simon and flanker conditions (i.e., whether FS - 00 = (F0 - 00) + (S0 - 00)). Indeed, RTs for multi-source conflict were significantly longer than expected from the additive model (M diff = 23.39 ms, SD = 51.87, t(39) = 2.85, p = 0.007). Multi-source conflict trials also resulted in significantly more errors (M diff = 4.47%, SD = 6.14, t(39) = 4.60, p < 0.001) than expected from the sum of Simon and flanker-induced errors.

### 3.2 Brain activity in Simon, flanker and multi-source conflicts

In the first step, we identified brain regions involved in solving conflicts, separately for: Simon, flanker and multi-source conditions (Figure 2, Suppl. Tables S1-S3). When contrasted with no-conflict condition, each type of the conflict included in MSIT evoked increased activity in the posterior medial frontal cortex (pMFC). Increased activity was also observed in dorsolateral prefrontal cortex (dlPFC) and the core regions of dorsal attention network (DAN): intraparietal sulci (IPS), frontal eye fields (FEF), and ventral premotor regions (vPM). Activity in flanker and multi-source conditions showed broader spatial extent and comprised clusters in higher-order visual cortex (lateral occipital complex, LOC), anterior insulas (aIns), and subcortical regions: thalamus and caudate nuclei. The observed pattern of activation was bilateral, however, activity in DAN was stronger in the left hemisphere. A number of regions also deactivated bilaterally during interference trials. When compared to no-conflict condition, each type of the conflict showed decreased activity in regions of the default mode network (DMN): ventromedial (vmPFC), ventrolateral (vlPFC), and dorsomedial prefrontal cortex (dmPFC); posterior sector of cingulate cortex with precuneus (Prec), as well as in angular gyri (Ang) and anterior temporal regions (aTmp). Again, a decrease of activity in flanker and multi-source conditions showed broader spatial extent and comprised clusters in ventral occipitotemporal cortex (lingual gyri and hippocampal regions), cuneus, and sensorimotor cortex (SM, central and precentral sulci and paracentral lobule).

**Figure 2.**
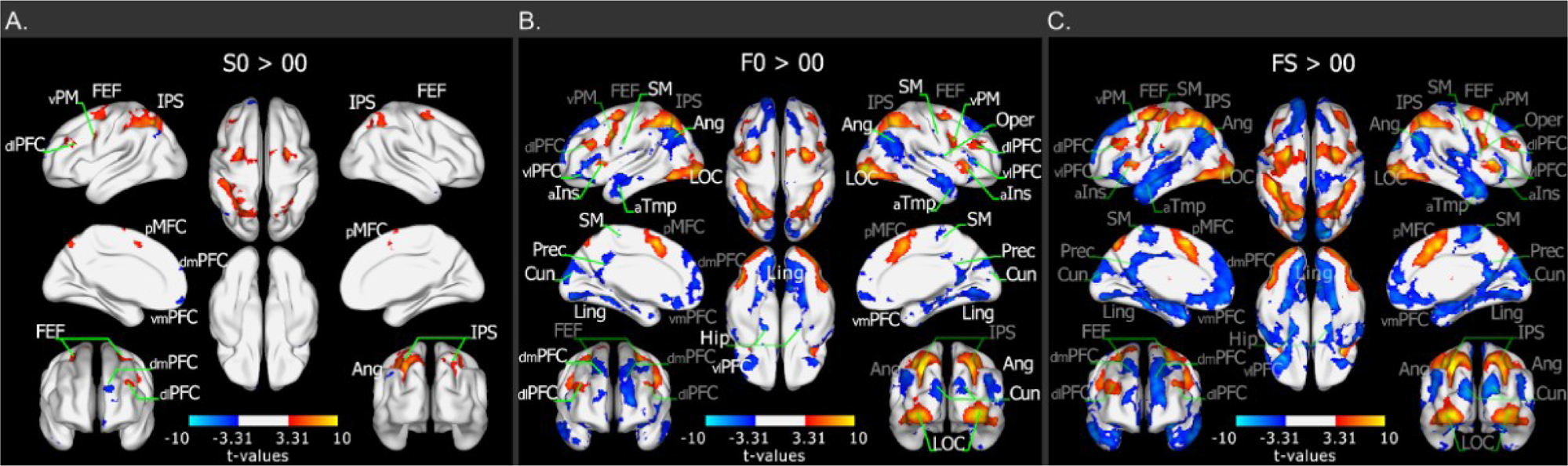
Brain regions involved in the three types of conflicts induced by MSIT. Contrasts between: A) Simon (S0) and no-conflict (00); B) flanker (F0) and no-conflict (00); C) multi-source (FS) and no-conflict (00) conditions. Contrast maps are thresholded at voxel-level p<0.001 and FWE-corrected (p<0.05) for cluster size. Labels written in white indicate first occurrence (from subplot A to C), whereas gray indicates that the label was already present in a previous subplot - such a presentation highlights the fact that no new brain regions were involved in FS conditions, despite the overall stronger/broader effects. For acronym explanation, see the list at the end of the manuscript.

For completeness, we also checked if some of the hemodynamic responses reflected the super-additive effects between Simon and flanker conflicts observed in behavioral performance. To this aim we performed a contrast [(FS + 00) vs (S0 + F0)] which effectively tests if the difference from no-conflict trials is the same for multi-source conflict as for the sum of Simon and flanker conditions (i.e., is equivalent to testing if FS - 00 = (F0 - 00) + (S0 - 00)). There were no significant clusters of super- or sub-additive activity in multi-source condition at the adopted significance level (i.e., cluster corrected at pFWE = 0.05).

### 3.3 Common and distinct regions involved by Simon and flanker conflicts

Conflict conditions included in the task showed substantial similarity of activation. To expose regions which showed a response both to Simon and flanker conflicts we performed a conjunction analysis of Simon and flanker-related differences from no-conflict conditions (Figure 3A, Suppl. Table S4). Conjunction was based on uncorrected maps, which were cut off at a lower voxel-level threshold of p < 0.01 (see Methods). The conjunction analysis confirmed that regions of common activation included pMFC, bilateral dlPFC and DAN nodes (IPS, FEF, vPM). Deactivation areas were also shown as largely common across conflict conditions, comprising bilateral regions of the DMN (vlPFC, vmPFC, dmPFC, Ang and aTemp), as well as the posterior part of the cingulate cortex and precuneus (Prec).

**Figure 3.**
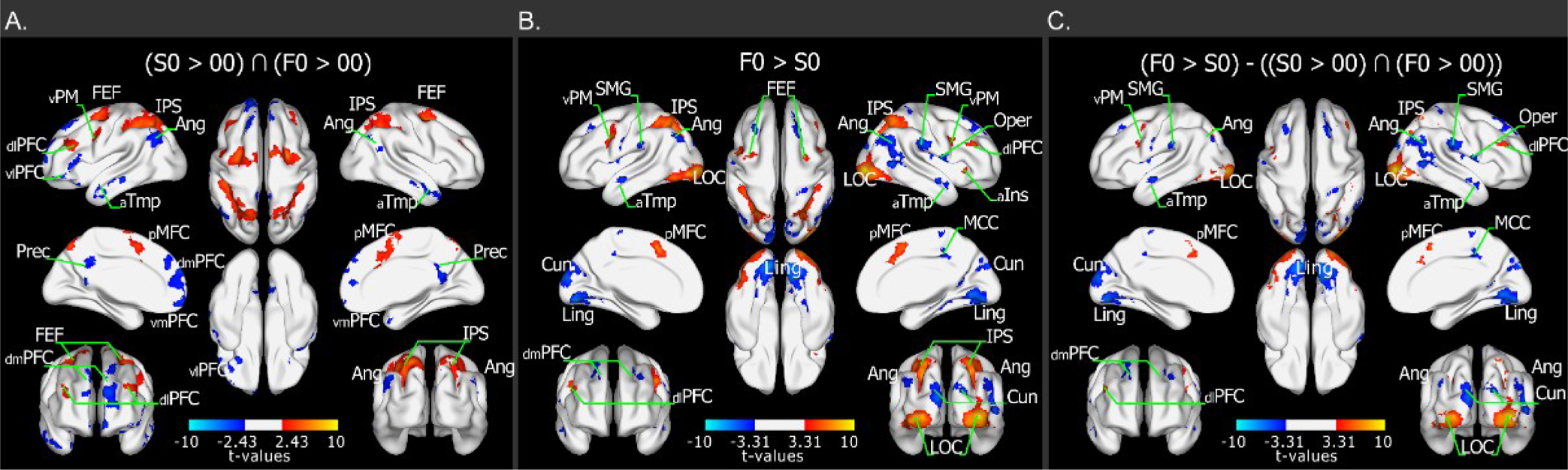
Common and distinct brain regions involved in resolving Simon and flanker conflicts. A) Conjunction of flanker (F0) and Simon (S0) effects. B) Difference between flanker- and Simon-related effects. C) Difference between flanker- and Simon-related effects masked by their conjunction map. A conjunction map was created with the voxel-level threshold p<0.01 used to encompass larger areas of common activity. Contrast maps in subplots B and C are thresholded at voxel-level p<0.001 and FWE-corrected (p<0.05) for cluster size. For acronym explanation, see the list at the end of the manuscript.

To identify the regions which show significantly different activity in Simon and flanker conflicts, we directly contrasted these two conditions. This comparison revealed that most of the regions activated by both Simon and flanker processing were also significantly more active in flanker than Simon conflicts (Figure 3B, Suppl. Table S5).

Regions identified by conjunction analysis were subsequently excluded from the F0 vs S0 contrast map to expose activity, which may be deemed specific for particular conflict (Figure 3C, Suppl. Table S6). The procedure indicated two classes of residual, conflict-specific activations: clusters directly adjacent to, or spatially separated from, the regions identified in conjunction analysis. The former confirmed that flanker conflict recruited broader areas of pMFC and DAN and deactivated broader areas of DMN. The latter proved pronounced differences between Simon- and flanker-related activity in the visual areas. Higher-order visual cortex (LOC) demonstrated significantly higher activity in the flanker compared to Simon condition, while other areas, such as parts of lingual gyrus and cuneus, showed significantly lower activity in flanker condition. The applied procedures also exposed significantly lower flanker-than Simon-related activity in the angular gyri and the bilateral supramarginal gyri (SMG). The latter effect was not earlier captured by contrasts with no-conflict condition.

### 3.4 Activity exceeding the time-on-task effects in Simon, flanker and multi-source conflicts

In line with the hypothesis that activations observed during conflict resolution reflects time-on-task effects, activity observed in MSIT showed progression of intensity and extent coinciding with increasing reaction time (i.e., Simon < flanker < multi-source conflict). The RT-related BOLD modulation was not identical across brain regions in different task conditions (Suppl. Fig. S1 and S2), which highlights the complex relationship between the task demands and modulation by time spent on task resolution. In other words, RT modulation may reflect conflict-independent time-on-task effect but also a conflict-specific RT variation. To de-confound activity related to conflict resolution from the effects of prolonged time spent on task, we modeled the activity in Simon, flanker and multi-source conditions, as expected solely on the basis of RT in co-conflict (00) condition and its RT modulation (00_RTmod_, see Methods 2.6.3) and compared them to observed activity.

Whole brain maps (Figure 4, Suppl. Figure S3, Suppl. Tables S7-9) revealed that in the Simon conflict, activity exceeding time-on-task effects was identified in left IPS. In flanker and multi-source conflicts higher-than-predicted activity included the pMFC, and additionally the right occipital pole (LOC) in flanker conflict. In addition to previously identified regions, comparison between the observed and RT-predicted signal in flanker and multi-source conditions showed higher-than-predicted activity in subcortical regions (caudate and putamen, see also Suppl. Figure S4) and white matter. Flanker and multi-source conditions also showed lower-than-predicted activity in visual cortex (calcarine sulcus, cuneus, fusiform gyri), in regions which were not previously found to engage in the task.

**Figure 4.**
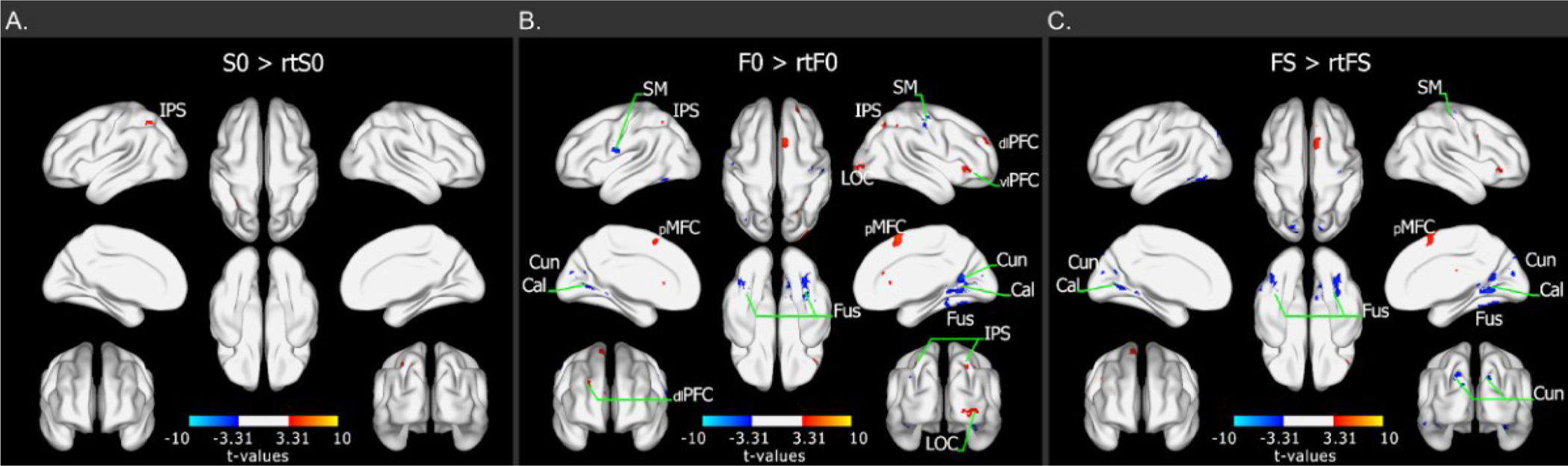
Regions with activations/deactivations exceeding the time-on-task effects in the three types of conflicts induced by MSIT. A) Contrast between the observed (S0) and RT-predicted (rtS0) activity in Simon conflict; B) the observed (F0) and RT-predicted (rtF0) activity in flanker conflict; C) the observed (FS) and RT-predicted (rtFS) activity in multi-source conflict. Contrast maps are thresholded at voxel-level p<0.001 and FWE-corrected (p<0.05) for cluster size. (The same contrasts are presented at p < 0.05 threshold in Suppl. Figure S3). For acronym explanation, see the list at the end of the manuscript.

### 3.5 Common and distinct regions exceeding the time-on-task effects in Simon and flanker conflicts

Finally, we repeated the conjunction analysis and direct comparison between Simon and flanker conflicts, as well as the masking procedure for those signals that exceed the time-on-task effects. The conjunction analysis confirmed that there were clusters of common above-time-on-task activity in the pMFC (encompassing dorsal anterior cingulate cortex [dACC] and pre-supplementary motor area [pre-SMA]) in both Simon and flanker conflicts. A similar pattern was also present in right aIns and right IFG (Figure 5A, Suppl. Table S10). It also confirmed common higher-than-predicted activity in right caudate and putamen, as well as in white matter across conflict conditions.

**Figure 5.**
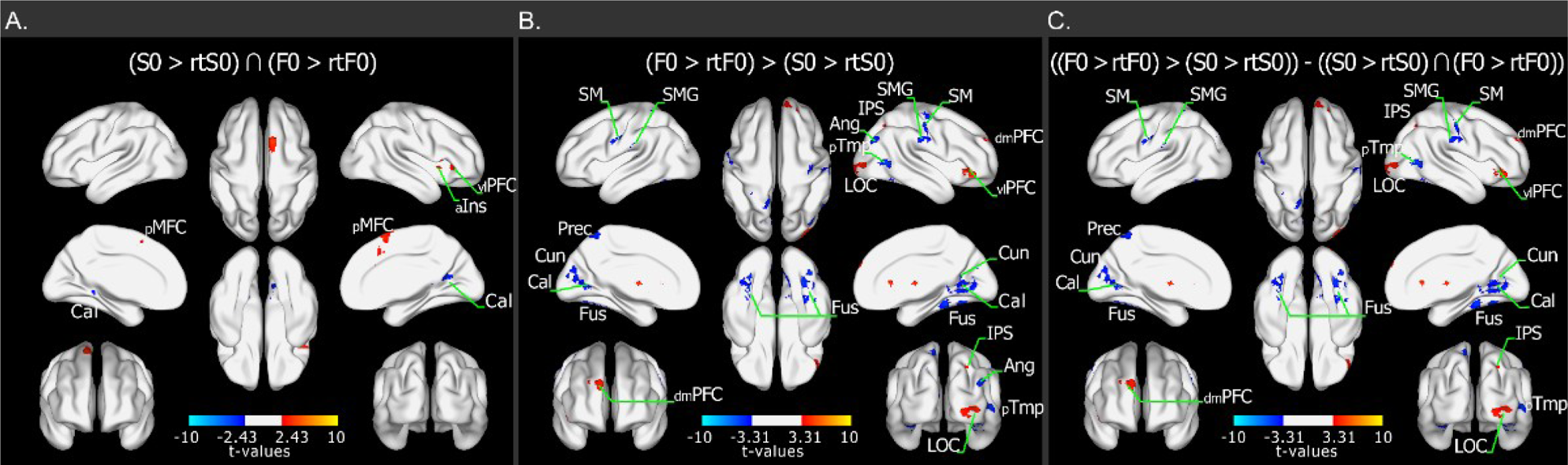
Common and distinct regions exceeding the time-on-task effects in Simon and flanker conflicts. A) Conjunction of activity exceeding the time-on-task effects in both Simon and flanker conflict conditions. B) Difference between activity exceeding the time-on-task effects in flanker and Simon conditions. For alternative analysis within RT-matched trials see Suppl. Fig. 5. C) Difference between activity exceeding the time-on-task effects in flanker and Simon conditions, masked by their conjunction. A conjunction map was created with the voxel-level threshold p<0.01 used to encompass larger areas of common activity. Contrast maps in subplots B and C are thresholded at voxel-level p<0.001 and FWE-corrected (p<0.05) for cluster size. For acronym explanation, see the list at the end of the manuscript.

The contrast analysis showed that activity in SMG, right sensorimotor cortex, occipital poles, calcarine sulcus, cuneus and fusiform gyri, deviated from RT-predicted signal significantly more in flanker than in Simon condition (Figure 5B, Suppl. Table S11; similar regions could also be observed in flanker vs Simon conflict comparison within RT-matched trials, Suppl. Fig. 5). All of these regions fell outside the conjunction map, as confirmed by the masking procedure (Figure 5C, Suppl. Table S12).

### 3.6 Time-on-task effects in pMFC

Additional analyses of pMFC ROIs enabled direct comparison with previous results describing the contribution of time-on-task effects within this area (Carp et al. 2010; Weissman and Carp 2013). These previous reports used a version of the task comprising no-conflict and multi-source conflict conditions; thus, we compared observed and RT-predicted signals evoked by multi-source condition (FS) within ROIs centered at the pMFC coordinates reported in these studies ([-3, 18, 51], Carp et al. 2010; [-6, 12, 49], Weissman and Carp, 2013; Figure 6A). We also analyzed the scope of the time-on-task effect across all the conflict conditions in ROIs defined within our study, in the respective main contrast between the conflict and no-conflict condition ([-2, 8, 58], S0 > 00; [-4, 10, 54], F0 > 00; [2, 8, 52], FS > 00; Figure 6A). The comparison between RT-predicted and observed activity in the pMFC ROIs showed that conflict-related activity exceeded time-on-task effects regardless of the task and specific coordinates employed for ROI definition (Figure 6B): the signal was significantly higher than predicted in ROIs defined by Carp et al. (2010) and Weissman and Carp (2013) for multi-source conflict, and, although non-significantly, also within ROI defined in our study; single-conflicts introduced in our task also evoked significantly higher activity than predicted on the basis of RT. Even though multi-source condition evoked the highest conflict, it also seemed less reliable in proving the difference between the expected and the observed responses (Figure 6B and C). Multi-source conflict condition may be less powerful to capture differences in question than uni-source conflicts if the latter evoke smaller but presumably less variable responses. We thus performed an additional simulation to check how the observed variability in the strength of the effect would translate into results of a smaller-sample study (N = 24; for comparison to the previous studies, N = 21, Carp et al. 2010; N = 24, Weissman and Carp, 2013). Random subsampling (n = 1000) showed that pairwise comparison between RT-predicted and observed signals in multi-source condition had only ∼13% chance of demonstrating the result below the conventional significance threshold (Figure 6C) in pMFC ROI defined in our data, and ≤ 68% chance in ROIs located at coordinates from previous studies. Interestingly, in single conflict conditions this probability was much higher (>83%). Thus, our complementary analysis confirms conflict-related signals in orthogonally defined pMFC ROIs, but also suggests that single conflicts may provide more robust effects than multi-source conflict condition.

**Figure 6.**
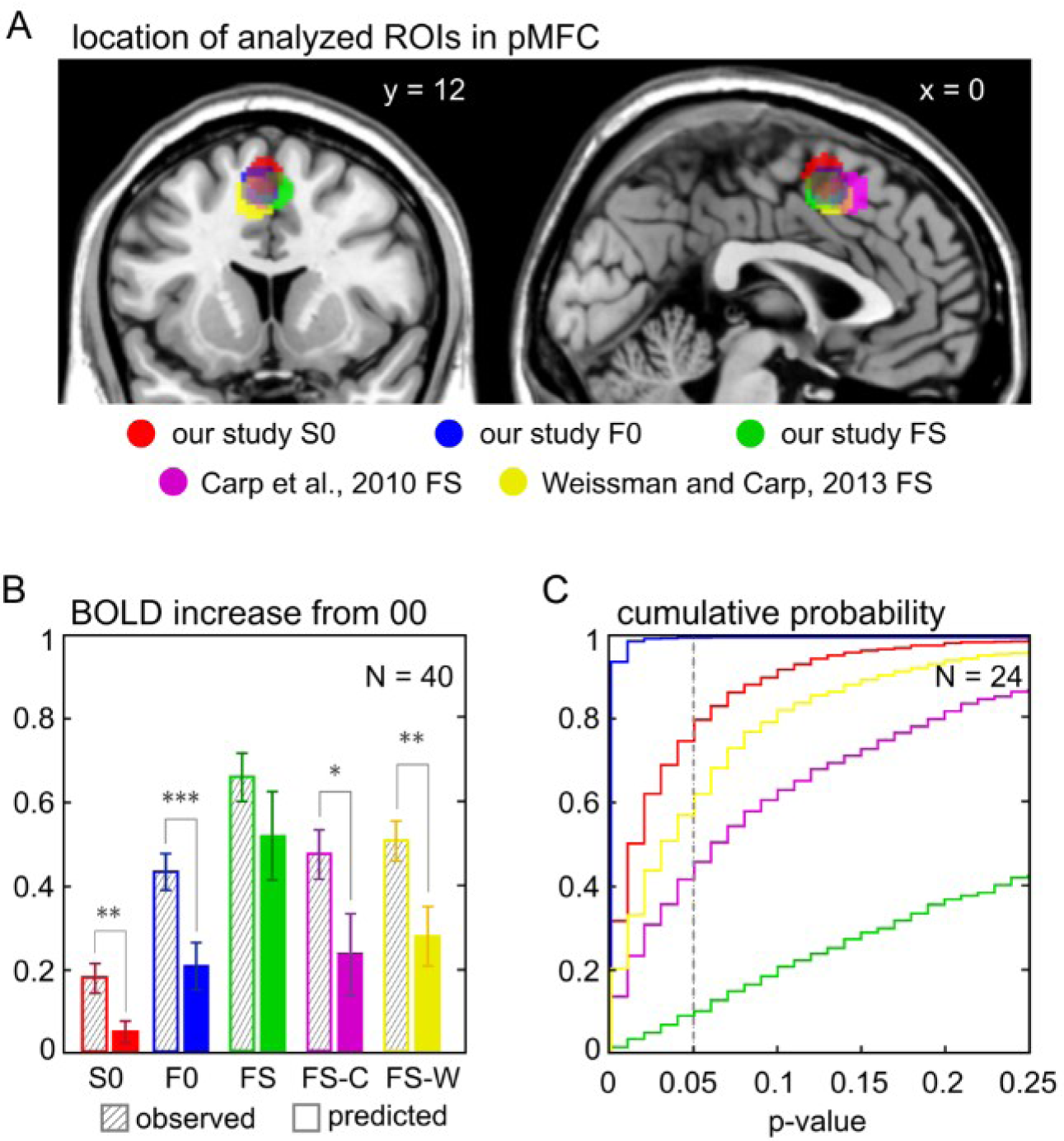
Time-on-task effects in pMFC. A) pMFC ROIs centered at the peak activity identified in multi-source conflict condition (FS > 00) in Carp et al. 2010 ([-3, 18, 51], magenta) and Weissman and Carp, 2013 ([-6, 12, 49], yellow), and in each conflict condition vs no-conflict in our study (S0 > 00, red, [-2, 8, 58]; F0 > 00, blue, [-4, 10, 54]; FS > 00, green, [2, 8, 52]). B) Comparison of RT-predicted and actual increase of activity (arbitrary units) in ROIs indicated in panel (A). Analysis on full group of N = 40, *** p < 0.001, ** p < 0.01, * p < 0.05. Outline colors code as in panel A, with grating marking real data and bars filled with a color indicating predicted values. C) Probability of observing a particular p-value when comparing RT-predicted and actual activity in pMFC ROIs indicated in panel A when the sample size equals N = 24 (simulation based on the data obtained in the present study, subsampled and tested 1000 times). Note that for N = 24 probability of reaching p = 0.05 threshold is higher (≥ 0.75) for single conflict conditions (S0, F0) and lower for FS condition (< 0.6) at each of the tested ROI locations. Color code as in panels A and B.

## 4. Discussion

Our study sought to determine shared and distinct correlates of conflict processing evoked by stimulus-response and stimulus-stimulus interference. The use of the extended MSIT (Sheth et al. 2012), the cognitive interference task which incorporates the Simon (stimulus-response) and flanker (stimulus-stimulus) conflicts, both separately and in combination, allowed us to compare conflict-related activity within a uniform design and set of stimuli. A number of regions showed common involvement in conflicts entailed by the MSIT, mainly within the dorsal attention network and medial prefrontal regions. Extensive common deactivation was also observed within the default mode network. In agreement with previous studies, which noted divergent activity depending on the type of interference, conflict-specific activity in the MSIT emerged mostly in the sensory/sensorimotor cortices (Egner and Hirsch 2005; Egner et al. 2007). Importantly, we were able to show that a part of the activity cannot be explained by time-on-task effects; neither within the regions engaged by a single type of interference nor within those involved across different conflicts.

Our results reinforce the idea that cognitive conflict engages additional mechanisms of cognitive control beyond those involved in less demanding, more automatized responses (Shenhav et al. 2013). This conclusion has been questioned by a series of studies which did not identify any BOLD activity over and above the effects explained by a longer time spent on processing the task in the presence of conflict (Carp et al. 2010; Grinband et al. 2011; Weissman and Carp 2013). These neuroimaging studies suggested that either there is no specific modulation of activity attributable to conflict occurrence or such modulation cannot be captured with the fMRI technique. In contrast, our findings support the coherence between fMRI and other experimental techniques that demonstrate conflict-dependent modulation (Ebitz and Platt 2015; Ebitz et al. 2020; Hanslmayr et al. 2008; Nigbur et al. 2011; Nigbur et al. 2012; Michelet et al. 2016).

### 4.1 Regions of common activity for both Simon- and flanker-type conflicts

Isolation of MSIT component conflicts revealed substantial similarity between Simon- and flanker-induced activation maps. Notably, a major part of this activity clearly overlapped with the large-scale intrinsic connectivity networks, i.e. networks stably identified with functional connectivity methods (Krienen et al. 2014; Power et al. 2011; Yeo et al. 2011). Regions of common activity for both Simon- and flanker-type conflicts were largely confined to the DAN (bilateral FEF, vPM and IPS/superior parietal lobule) and pMFC (dACC and pre-SMA), a major part of the cingulo-opercular network (CON). With regards to deactivation, we observed a down-modulation of the DMN (dmPFC, bilateral angular gyri and anterior temporal cortex). The regions of the DAN are known to coordinate action and spatial relationship between the objects and the effectors (eyes/hands; Andersen and Buneo 2002; Colby and Goldberg 1999). The DAN is also the main network implicated in attention, both in terms of directing attention to a specific location and to specific properties of stimuli (Corbetta and Shulman 2002; Squire et al. 2013). Increased activity clearly delineating the DAN may reflect the spatial nature of both Simon and flanker tasks. Even though stimulus-response mapping is congruent in flanker, both tasks rely on selection between spatially distributed elements. The inherently spatial character of Simon and flanker tasks can be explained via a comparison with the Stroop task, in which stimuli features (color and meaning) are bound to the same object (word) and thus do not entail spatial processing within the task execution (Stroop 1935). Thus, activity within the DAN likely reflects directing attention to, and selecting between, the MSIT stimuli. Increased activity within pMFC has been repetitively found in conflict tasks and in other contexts related to action selection such as reward and error monitoring, reacting to surprise, regulating arousal etc. (Ebitz and Hayden 2016; Kolling et al. 2016; Heilbronner and Hayden 2016; Ullsperger et al. 2014). Most broadly, pMFC has been proposed to mediate the interaction between motivational state and motor plans. From the network perspective, activity within CON has been shown to increase in phase at the initiation and completion of a particular activity and remain at elevated levels throughout the activity (Dosenbach et al. 2006). This suggested that CON has an important contribution to the implementation and maintenance of a specific activity rule. Much less is known about how deactivation of DMN may contribute to the execution of effortful tasks. This is due to the bias within the neuroimaging studies that prevalently report the one-sided contrasts (Cieslik et al. 2015, but see McDonald et al. 2017). However, we hypothesize that increased activity in the DAN and decreased activity in the DMN reflect the two sides of the same coin, supporting a maintained, elevated focus on external stimuli. Anticorrelation of the DAN and pMFC on the one side and the DMN on the other has been observed even in the absence of a task (Fox et al. 2005; He et al. 2007; Jurewicz et al. 2020). Work on the functional architecture of the brain suggest a hierarchy of the networks that form the extrinsic and intrinsic systems: the former related to taking action in the environment (encompassing the DAN), and the latter related to processes disengaged from external stimulation (encompassing the DMN; Golland et al. 2008; Margulies et al. 2016). Thus, the parallel increase and decrease of activity in large-scale networks involved in MSIT is probably one of the basic mechanisms that orchestrate brain resources to effectively support the visuomotor task and externally driven action.

Importantly, activations and deactivations in the DAN and the DMN were gaining area and strength, in an order consistent with the progression of RTs (from Simon to flanker and multi-source conflicts). The processes which closely correlate with the reaction times are collectively referred to as the “time-on-task effects’’. This set includes the mechanisms of attention, movement preparation etc., and plays an essential role in execution of any task. Examination of RT-related effects in our data revealed that indeed time-on-task effects could explain a large portion of DAN and DMN activations and deactivations, indicating that substantial part of the activity within these two networks is non-specific to conflict occurrence. (Activity exceeding time-on-task effects was present in IPS in both Simon and flanker conflicts, but it was absent from the conjunction of the two activity maps, indicating spatially non-overlapping conflict-related activity in Simon and flanker - see the next section. Weaker effects could also be observed across DAN regions after lowering the voxel-level significance threshold [Suppl. Figure S3]). A notable exception among the regions commonly activated by all the tasks was pMFC. Although some part of the activity in pMFC was explained by the time-on-task, parts of pMFC exhibited an up-modulation exceeding RT-based predictions (Figs. 5 and 6, and Suppl. Figs. 3 and 5; see the Discussion section 4.4).

### 4.2 Conflict-specific activity

Conflict-specific effects were identified predominantly for the flanker task: there were flanker-specific increases of the signal within the regions commonly involved in MSIT conditions and flanker-specific activations as well as deactivations in other parts of the brain. Most notably, flanker-type interference caused widespread increases and decreases of the signal in several regions of the visual cortex. Occipital poles, extending to inferior and superior occipital gyri and encompassing the lateral occipital complex (LOC) were activated only by the flanker conflict. As predicted for the conflict-specific response, a part of the activity within these regions could not be explained by simple time-on-task effects. Involvement of these regions, known to take part in the analysis of letters, symbols, and digits, likely reflected enhanced processing of MSIT stimuli in the presence of flanker interference. At the same time, parts of lingual and fusiform gyri on the basal surface of the brain, as well as parts of cuneus, were deactivated by flanker conflict. The retinotopic mapping of this part of the visual cortex suggests that processing of the task-irrelevant peripheral space may be suppressed during stimulus-stimulus conflict resolution (Wu et al. 2012; see Fan et al. 2007 for a similar observation). Calcarine sulcus, as well as parts of fusiform gyri were not significantly up- or down-modulated when compared to no-conflict condition, even though RT-related variability in these regions predicted an increase of the signal with longer RT. Possibly, the activity in calcarine and fusiform gyri may have saturated quickly, or such an increase was inhibited in conflict conditions. Together, a complex pattern of flanker conflict-related changes in the visual cortex suggested a multifaceted enhancement of the task-relevant information (cf. Wiesman and Wilson 2020). Additionally, activity exceeding time-on-task effects was present in IPS in both Simon and flanker conflicts in spatially non-overlapping regions. Our results support the view that response conflict in flanker condition influences the whole path of information processing extending back to the visual cortex over the period when the task is solved. This observation expands the idea of “stimulus biasing”, discussed as a mechanism for conflict resolution (Botvinick et al. 2001; Cohen et al. 1990; Egner and Hirsch 2005), which consists in excitatory biasing of task-relevant feature. MSIT does not allow for anticipatory preparation of the relevant feature or dimension of stimuli – a target can only be selected when the whole stimulus has been integrated. Modulation of activity at different levels of the information processing pathway points at the distributed nature of action selection (Cisek and Kalaska 2010). In this task, modulation of the visual cortex appears to play an active role in decision-making and response selection via top-down modulation and feedback.

Flanker-specific deactivation was also observed in some other regions, including SMG (Figure 3C and Figure 5C). Subtractive logic of BOLD analysis makes it ambiguous whether lower Simon-than flanker-related activity means a flanker-specific deactivation or a Simon-specific activation (Figure 3C). However, RT-based comparisons help disambiguate this observation: activity within SMG in flanker condition was lower than predicted on the basis of RT and it was so to a greater extent than in Simon condition (Figure 5C). Thus, modulation within SMG reflects another example of a flanker-related suppression of hemodynamic response, rather than Simon-specific increase. Spatial localization of this effect agrees with conflict-specific effects identified earlier in a similar comparison of Simon and Stroop tasks (Egner et al. 2007). We may speculate that suppression in this region may prevent premature responses in flanker conditions, enabling longer analysis of the stimuli in this more demanding task (Simos et al. 2017). Lack of Simon-specific changes of activity beyond conflict-dependent enhancement in IPS may be related to the fact that the mapping between the spatial position of a target and the required response is computed within the DAN. Consequently, stimulus-response mapping, targeted by Simon-type interference, depends on the neural circuits that overlap with regions supporting response selection, attention orienting and maintenance etc., recruited by all task conditions.

### 4.3 Interaction between Simon- and flanker-type conflicts

Resources shared between the tasks could also have been revealed in the analysis of the interaction of Simon and flanker conflicts. Behavioral performance in multi-source interference condition was worse than expected from adding the influences of Simon and flanker interference. This result is consistent with previous findings on extended MSIT (Wiesman and Wilson 2020) and other tasks combining Simon and flanker conflicts (Fan et al. 2003; Frühholz et al. 2011; Hommel 1997; Mückschel et al. 2016) and may indicate shared processing mechanisms between the tasks (Egner 2008). However, despite superadditive RTs and error rates in multi-source condition, we did not observe any reliable interactive effects in the BOLD signal. Extensive discussion of potential sources of diverging behavioral and neural results in multi-source conflict task can be found in Rey-Mermet et al. 2019. One possible explanation is that the saturation of the hemodynamic response, which was boosted in the highly demanding multi-source condition, left no space for superadditive effects (Schei et al. 2011). It is also possible that the interaction influenced the dynamics, not the overall magnitude of neuronal responses, and the BOLD signal could not detect such differences. Indeed, some superadditive effects between Simon and flanker conflicts in multi-source condition have been previously detected, and localized in the parietal and premotor cortex, in MEG source-reconstructed signals of the selected oscillation bands (Wiesman and Wilson 2020; Wiesman et al. 2020). The ERP analysis of the EEG signals obtained within the same task (Dzianok et al. 2022) may help to resolve and explain discrepancies between behavioral and fMRI results.

### 4.4 The scope of time-on-task effects

The question about the role of conflict in pMFC function is particularly vital as this region has been frequently implicated in cognitive control and targeted in related research (Botvinick et al. 1999; Shenhav et al. 2013; Ullsperger and von Cramon 2001). It is important to note that while we found modulation of the BOLD activity attributable to cognitive conflict in pMFC, that does not mean that we identified an abstract “conflict” signal, postulated in some accounts on cognitive control (Shenhav et al. 2013). Our results are consistent with multiple alternative views on the mechanisms of conflict resolution. While conflict-dependent signals may relate to conflict monitoring, they may also result from multiple competing, simultaneously available actions, or increased biasing of stimuli and/or responses (Cieslik et al. 2015; Ebitz et al. 2020). The pMFC encompasses the pre-supplementary motor area (pre-SMA) and anterior midcingulate cortex (aMCC) or, more often invoked, dorsal anterior cingulate cortex (dACC). While pre-SMA has been more strongly associated with control of motor output, e.g., selecting the appropriate response between different response alternatives, aMCC has been primarily associated with enhancing behavioral plans relevant to task demands against competing behavioral alternatives. However, both of these regions have been found jointly involved in many tasks, and their clear functional dissociation is rather difficult (Nachev et al. 2008). Similarly, in our task, we observed distributed activity in both Simon and flanker conflicts spanning dACC and pre-SMA. In both tasks, the peak of activity and conflict-dependent effects were localized predominantly in pre-SMA (cf. Beldzik et al. 2015, Beldzik and Ullsperger 2023). This finding is similar to the previous report on the mechanisms for resolving stimulus-stimulus and stimulus-response interference, which showed the region of common activity in pre-SMA and stimulus-stimulus specific activity in dACC (Li et al. 2017). However, on the basis of our results, we cannot conclude whether Simon and flanker-related processing is localized to discrete neuronal populations in the pMFC. These questions should be further addressed with other experimental techniques. Indeed, a recent study using single-cell recordings in human dACC suggested that the populations of cells significantly modulated by Simon and flanker conflicts were almost entirely non-overlapping (Ebitz et al. 2020).

Significant differences obtained in our study between RT-predicted and conflict-induced activations contrast with the results of similar analyses performed in previous research (Carp et al. 2010; Grinband et al. 2011; Weissman and Carp 2013). Importantly, studies by Carp and colleagues which showed that activity in pMFC is fully explained by time-on-task effects also used MSIT, although in a classic version with only one, multi-source conflict condition. Thus, divergent imaging outcomes cannot stem from task-specific features of neural processing, but rather some experimental details that had a decisive impact upon the results. Current and previous studies differed in multiple aspects of experimental settings, such as methods of data acquisition, distribution and timing of the stimuli, number of regressors etc. Although we cannot conclude which factor is crucial for explaining the divergent results, we want to consider yet another factor–the power of statistical analyses, as playing an important role here. A priori power analysis in neuroimaging research is extremely difficult due to the huge variability of both experimental designs and cognitive processes under study (Mumford 2012). However, a meta-research on the power of fMRI analyses suggests that studies based on samples of around 20 subjects are only able to capture relatively large effect sizes (Button et al. 2013; Poldrack et al. 2017). Our study was based on a larger sample (n = 40), which almost doubled the samples used in previous studies. An additional test performed for ROIs, located at the peaks of pMFC activity in multi-source conflict condition reported by Carp et al. (2010) and by Weissman and Carp (2013), performed for our full (n = 40) dataset, indicated significant overshoot of observed signal over the time-on-task prediction. On the other hand, the predicted difference was not large enough in FS condition to reach significance within the ROI indicated in our study. When our sample was randomly limited to N = 24, probability that the comparison between RT-predicted and actual activity was below the conventional significance threshold (p < 0.05) was lower for all FS-based ROIs. The same analysis performed for S0 and F0 conditions revealed considerably stronger conflict-specific effects. This suggests that a strong, cumulative conflict may not always be the most apt to study the time-on-task vs. conflict-related effects. Signs of conflict-related signals exceeding time-on-task effects were already reported by Carp et al. (2010) in the whole brain analysis, in bilateral parietal cortices. Within these regions, higher activity was found in multi-source conflict condition in comparison to no-conflict condition when trials were subsampled to have matching RT. Several brain regions were also shown to activate beyond the time-on-task effects in a different conflict study, which tested oculomotor reactions in an anti-saccade task (Beldzik et al. 2015) and Stroop and Simon tasks (preprint Beldzik & Ullsperger, 2023). Activated maps included frontal and parietal areas as well as pMFC (in particular the pre-SMA), further supporting our finding that these regions engage in different conflicts and show higher activity than expected simply on the basis of RT.

### 4.5 Conclusions

Conflict-induced changes in brain activity were predominantly present in common regions of DAN, pMFC and DMN regardless of the type of conflict; with their strength and extent scaling from faster to slower-resolved tasks. Their coordinated activity suggested a joint support for externally driven, visuo-spatial tasks, not limited to conflict resolution. However, despite the uniform type of stimuli used for different conditions, conflict-specific activations were also present in the sensory/sensorimotor cortices. Thus, sensory resources were differentially recruited during various tasks, supporting the idea of distributed mechanisms for resolving conflicts. Although time-on-task effects did explain a large part of observed activations, the scope of this effect was not exhaustive, i.e., several regions exhibited the activation above and below the levels predicted on the basis of RT. In particular, activity within pMFC (encompassing dACC and pre-SMA) could not be reduced to the prolonged time spent on task during resolution of different types of conflicts. Our study provides new evidence that, with increased statistical power, conflict-related modulation of the BOLD signal can be observed in regions associated with cognitive control. This insight helps to explain conflicting findings in the field of cognitive control studies and validates neuroimaging research on conflict processing, highlighting the need for the increased sensitivity for capturing weaker–but critical–cognitive effects.

## Declarations of Competing Interest

Authors declare that they have no conflict of interest.

## CRediT authorship contribution statement

JW: Conceptualization, Investigation, Methodology, Data curation, Formal analysis, Visualization, Writing (review & editing); KJ: Conceptualization, Methodology, Formal analysis, Visualization, Writing (original draft), Writing (review & editing), Supervision; PD: Conceptualization, Investigation, Data curation, Formal analysis, Writing (review & editing); IA: Conceptualization, Writing (review & editing); KP: Conceptualization, Writing (review & editing); TW: Conceptualization, Methodology, Writing (review & editing), Funding acquisition, Supervision; EK: Conceptualization, Visualization, Writing (review & editing), Funding acquisition, Supervision

## Data and code availability

The unthresholded and unmasked group-level whole-brain results maps are available at Neurovault repository: https://neurovault.org/collections/NTRDGLJW/

All analysis codes are available upon request.

## Acknowledgements

This work was supported by the National Science Centre, Poland; grant number 2016/20/W/NZ4/00354. We thank Ms Alicja Dobrzykowska for her help in organizing MRI data acquisition.

## Abbreviations

FEF: frontal eye field
IPS: intraparietal sulcus
vPM: ventral premotor cortex
Ang: angular gyrus
aTmp: anterior temporal gyrus
pTmp: posterior temporal gyrus
LOC: lateral occipital complex
aIns: anterior insula
Thal: thalamus
Caud: caudate
Ling: lingual gyrus
Cun: cuneus
Prec: precuneus
Fus: fusiform gyrus
Hip: hippocampus
Oper: operculum
pMFC: posterior medial frontal cortex
mPFC: medial prefrontal cortex
dlPFC: dorsolateral prefrontal cortex
dmPFC: dorsomedial prefrontal cortex
vmPFC: ventromedial prefrontal cortex
vlPFC: ventrolateral prefrontal cortex
SM: sensorimotor cortex
SMA: supplementary motor area
pre-SMA: pre-supplementary sensorimotor area
cal: calcarine sulcus
SMG: supra marginal gyrus
MCC: midcingulate cortex
aMCC: anterior midcingulate cortex
dACC: dorsal anterior cingulate cortex
wm: white matter
Put: putamen
DMN: default-mode network
DAN: dorsal attention network
MSIT: multi-source interference task
FS: Simon and flanker effects combined in one condition
F0: flanker effect condition
S0: Simon effect condition
00: no-conflict condition

## Notes

Funding: this work was supported by the National Science Centre, Poland; grant number 2016/20/W/NZ4/00354.

### Competing Interest Statement

The authors have declared no competing interest.

### Summary of Updates

In the revised version of the manuscript we clarify and illustrate (in supplementary figures) the reasoning related to the chosen design of fMRI analyses.

