## supplementary tables for "Common and distinct BOLD correlates of Simon and flanker conflicts which can(not) be reduced to time-on-task effects"

### Supplementary materials include:

1) Supplementary figures S1-S5

2) Supplementary tables S1-S12, included in this file

3) Unthresholded and unmasked group-level whole-brain results maps are available at Neurovault repository: https://neurovault.org/collections/NTRDGLJW/.

Table S1

Regions differing in BOLD activity between Simon vs no-conflict conditions.

| **Coordinates MNI** | | | **Cluster size** | **(% of cluster) AAL Region** | **Cluster pFWE** | **Cluster pFDR** | **Cluster p(uncor)** | **Peak T** | **Cluster label** |
|---|---|---|---|---|---|---|---|---|---|
| **x** | **y** | **z** | | | | | | | |
| **S0 > 00** | | | | | | | | | |
| -26 | -60 | 48 | 2166 | (38%) Parietal_Inf_L<br>(32%) Parietal_Sup_L<br>(8%) Postcentral_L<br>(8%) OUTSIDE<br>(7%) Occipital_Mid_L<br>(5%) Precuneus_L | < 0.001 | < 0.001 | < 0.001 | 9,44 | **IPS L** |
| -28 | 2 | 60 | 958 | (47%) Frontal_Sup_2_L<br>(22%) Precentral_L<br>(15%) Frontal_Mid_2_L<br>(13%) OUTSIDE | < 0.001 | < 0.001 | < 0.001 | 6,53 | **FEF L** |
| 26 | 2 | 54 | 551 | (70%) Frontal_Sup_2_R<br>(15%) OUTSIDE<br>(9%) Frontal_Mid_2_R<br>(7%) Precentral_R | < 0.001 | < 0.001 | < 0.001 | 6,39 | **FEF R** |
| -2 | 8 | 58 | 390 | (50%) Supp_Motor_Area_L<br>(39%) Supp_Motor_Area_R<br>(9%) Frontal_Sup_2_R | < 0.001 | < 0.001 | < 0.001 | 6,38 | **pMFC** |
| 32 | -52 | 46 | 669 | (43%) Parietal_Sup_R<br>(16%) Parietal_Inf_R<br>(12%) Occipital_Sup_R<br>(12%) OUTSIDE<br>(8%) Angular_R<br>(6%) Precuneus_R | < 0.001 | < 0.001 | < 0.001 | 5,57 | **IPS R** |
| -38 | 0 | 36 | 158 | (97%) Precentral_L | 0,02 | 0 | 0 | 4,64 | vPM |
| -36 | -54 | -28 | 152 | (71%) Cerebelum_6_L<br>(29%) Cerebelum_Crus1_L | 0,02 | 0 | 0 | 4,64 | |

| | | | | | | | | | |
|---|---|---|---|---|---|---|---|---|---|
| -32 | 40 | 30 | 184 | (67%) Frontal_Mid_2_L<br>(31%) Frontal_Inf_Tri_L | 0,01 | 0 | < 0.001 | 4,31 | **dlPFC** |
| **S0 < 00** | | | | | | | | | |
| -4 | 62 | -10 | 259 | (70%) Frontal_Med_Orb_L<br>(14%) Frontal_Med_Orb_R<br>(8%) Rectus_L | 0 | 0 | < 0.001 | 5,42 | vmPFC |
| -60 | -64 | 32 | 162 | (74%) Angular_L<br>(26%) OUTSIDE | 0,01 | 0,01 | < 0.001 | 5,3 | **Ang** |
| -14 | 62 | 24 | 167 | (71%) Frontal_Sup_2_L<br>(28%) Frontal_Sup_Medial_L | 0,01 | 0,01 | < 0.001 | 5,24 | dmPFC |
| 28 | 20 | -26 | 159 | (44%) Temporal_Pole_Mid_R<br>(37%) Temporal_Pole_Sup_R<br>(12%) OFCpost_R<br>(6%) OUTSIDE | 0,02 | 0,01 | 0 | 4,69 | **aTmp R** |

Functional regions are reported with corresponding MNI coordinates of peak activity within each cluster as well as cluster size. Additionally noted is the percentage of cluster's voxel in each anatomical region defined using the AAL3 atlas. Only regions with more than 5% of all cluster's voxel are noted. Results are thresholded at $p<0.001$ for voxel and FWE-corrected ($p<0.05$) for cluster size. Functional labels which are visible on the figures are marked in bold.

Table S2

Regions differing in BOLD activity between flanker vs no-conflict conditions.

| Coordinates MNI x | y | z | Cluster size | (% of cluster) AAL Region | Cluster pFWE | Cluster pFDR | Cluster p(uncor) | Peak T | Cluster label |
|---|---|---|---|---|---|---|---|---|---|
| **F0 > 00** | | | | | | | | | |
| -26 | -58 | 48 | 3837 | (31%) Parietal_Sup_L | < 0.001 | < 0.001 | < 0.001 | 12,12 | **IPS L** |
| | | | | (31%) Parietal_Inf_L | | | | | |
| | | | | (12%) OUTSIDE | | | | | |
| | | | | (9%) Postcentral_L | | | | | |
| | | | | (8%) Occipital_Mid_L | | | | | |
| | | | | (5%) Precuneus_L | | | | | |
| 32 | -90 | -2 | 11802 | (12%) OUTSIDE | < 0.001 | < 0.001 | < 0.001 | 11,7 | **LOC (R+L)** **IPS R** |
| | | | | (7%) Occipital_Inf_R | | | | | |
| | | | | (7%) Parietal_Sup_R | | | | | |
| | | | | (7%) Cerebelum_6_R | | | | | |
| | | | | (6%) Cerebelum_Crus1_L | | | | | |
| | | | | (6%) Cerebelum_6_L | | | | | |
| | | | | (6%) Occipital_Inf_L | | | | | |
| | | | | (5%) Occipital_Mid_R | | | | | |
| -30 | 0 | 60 | 6910 | (20%) Precentral_L | < 0.001 | < 0.001 | < 0.001 | 10,64 | **pMFC + FEF (R+L)** |
| | | | | (11%) OUTSIDE | | | | | |
| | | | | (11%) Supp_Motor_Area_L | | | | | |
| | | | | (10%) Supp_Motor_Area_R | | | | | |
| | | | | (9%) Frontal_Sup_2_R | | | | | |
| | | | | (6%) Frontal_Sup_2_L | | | | | |
| | | | | (5%) Precentral_R | | | | | |
| | | | | (5%) Frontal_Inf_Tri_L | | | | | |
| | | | | (5%) Frontal_Mid_2_L | | | | | |
| 32 | 28 | 2 | 616 | (65%) Insula_R | < 0.001 | < 0.001 | < 0.001 | 8,2 | **aIns R** |
| | | | | (15%) OUTSIDE | | | | | |
| | | | | (8%) Frontal_Inf_Oper_R | | | | | |
| | | | | (7%) Frontal_Inf_Tri_R | | | | | |

| | | | | | | | | | |
|---|---|---|---|---|---|---|---|---|---|
| -32 | 18 | 10 | 275 | (91%) Insula_L | < 0.001 | < 0.001 | < 0.001 | 6,57 | **aIns L** |
| -12 | -16 | 12 | 279 | (31%) Thal_VL_L | < 0.001 | < 0.001 | < 0.001 | 6,56 | Thal L<br>Caud L |
| | | | | (27%) OUTSIDE | | | | | |
| | | | | (10%) Caudate_L | | | | | |
| | | | | (8%) Thal_VA_L | | | | | |
| | | | | (7%) Thal_VPL_L | | | | | |
| | | | | (5%) Thal_LP_L | | | | | |
| 34 | 54 | 30 | 895 | (73%) Frontal_Mid_2_R | < 0.001 | < 0.001 | < 0.001 | 6,35 | **dlPFC** |
| | | | | (15%) OUTSIDE | | | | | |
| | | | | (6%) Frontal_Inf_Tri_R | | | | | |
| | | | | (6%) Frontal_Sup_2_R | | | | | |
| 16 | -24 | 14 | 309 | (33%) Caudate_R | < 0.001 | < 0.001 | < 0.001 | 5,82 | Thal R<br>Caud R |
| | | | | (20%) Thal_VL_R | | | | | |
| | | | | (20%) OUTSIDE | | | | | |
| | | | | (6%) Thal_LP_R | | | | | |
| | | | | (6%) Thal_PuM_R | | | | | |
| -24 | -72 | -50 | 125 | (60%) Cerebelum_8_L | 0,04 | 0 | 0 | 4,98 | |
| | | | | (38%) Cerebelum_7b_L | | | | | |
| **F0 < 00** | | | | | | | | | |
| 32 | -42 | -8 | 5788 | (16%) Lingual_R | < 0.001 | < 0.001 | < 0.001 | 8,11 | **Ling R** |
| | | | | (10%) Cuneus_R | | | | | **Cun (R+L)** |
| | | | | (9%) Cuneus_L | | | | | |
| | | | | (8%) ParaHippocampal_R | | | | | |
| | | | | (7%) Precuneus_R | | | | | **Prec** |
| | | | | (6%) Fusiform_R | | | | | |
| | | | | (5%) Calcarine_R | | | | | |
| | | | | (5%) Calcarine_L | | | | | |
| | | | | (5%) Occipital_Sup_L | | | | | |
| 54 | -72 | 38 | 2867 | (44%) Temporal_Mid_R | < 0.001 | < 0.001 | < 0.001 | 7,06 | **Ang R** |
| | | | | (29%) Angular_R | | | | | |
| | | | | (9%) OUTSIDE | | | | | |
| | | | | (8%) Occipital_Mid_R | | | | | |

| | | | | | | | | | |
|---|---|---|---|---|---|---|---|---|---|
| | | | | (7%) Temporal_Sup_R | | | | | |
| 54 | 32 | 6 | 347 | (99%) Frontal_Inf_Tri_R | < 0.001 | < 0.001 | < 0.001 | 7 | |
| -24 | -44 | -6 | 387 | (47%) Fusiform_L<br>(23%) ParaHippocampal_L<br>(18%) Lingual_L<br>(6%) Hippocampus_L<br>(5%) OUTSIDE | < 0.001 | < 0.001 | < 0.001 | 6,86 | Fus<br>+ Hip<br>+ **Ling** |
| 0 | 62 | -6 | 3623 | (31%) Frontal_Sup_2_L<br>(22%) Frontal_Sup_Medial_L<br>(11%) Frontal_Med_Orb_L<br>(10%) Frontal_Mid_2_L<br>(9%) Frontal_Med_Orb_R | < 0.001 | < 0.001 | < 0.001 | 6,81 | **vmPFC (R+L)**<br>**+ vlPFC (R+L)**<br>**+ dmPFC L** |
| -18 | -6 | -16 | 690 | (37%) Hippocampus_L<br>(27%) ParaHippocampal_L<br>(11%) Amygdala_L<br>(8%) OUTSIDE<br>(5%) Temporal_Pole_Sup_L | < 0.001 | < 0.001 | < 0.001 | 6,81 | **Hip L** |
| 36 | -18 | 16 | 545 | (54%) Rolandic_Oper_R<br>(22%) Insula_R<br>(9%) Postcentral_R<br>(7%) Temporal_Sup_R<br>(6%) OUTSIDE | < 0.001 | < 0.001 | < 0.001 | 6,75 | **Oper R** |
| 26 | 32 | 58 | 861 | (70%) Frontal_Sup_2_R<br>(16%) Frontal_Sup_Medial_R<br>(8%) OUTSIDE<br>(5%) Frontal_Mid_2_R | < 0.001 | < 0.001 | < 0.001 | 6,53 | **dmPFC R** |
| -46 | -66 | 42 | 2683 | (37%) OUTSIDE<br>(29%) Angular_L<br>(10%) Temporal_Mid_L<br>(8%) Occipital_Mid_L<br>(7%) Parietal_Inf_L | < 0.001 | < 0.001 | < 0.001 | 6,47 | **Ang L** |

| | | | | | | | | | |
|---|---|---|---|---|---|---|---|---|---|
| | | | | (6%) SupraMarginal_L | | | | | |
| 54 | -6 | -14 | 1202 | (49%) Temporal_Mid_R<br>(20%) Temporal_Pole_Mid_R<br>(17%) Temporal_Sup_R<br>(8%) Temporal_Pole_Sup_R | < 0.001 | < 0.001 | < 0.001 | 6,45 | **aTmp R** |
| -50 | 38 | 2 | 2404 | (36%) Temporal_Mid_L<br>(16%) Frontal_Inf_Tri_L<br>(14%) OUTSIDE<br>(8%) Frontal_Inf_Orb_2_L<br>(6%) OFCpost_L | < 0.001 | < 0.001 | < 0.001 | 6,31 | **aTmp L** |
| -10 | -86 | -10 | 692 | (79%) Lingual_L<br>(8%) Fusiform_L<br>(6%) Calcarine_L<br>(6%) Cerebelum_6_L | < 0.001 | < 0.001 | < 0.001 | 6,05 | **Ling L** |
| 40 | 30 | -14 | 162 | (51%) OFCpost_R<br>(20%) Frontal_Inf_Orb_2_R<br>(9%) OFCant_R<br>(9%) OFCmed_R | 0,01 | 0 | < 0.001 | 5,14 | **vlPFC** |
| 38 | -16 | 40 | 510 | (26%) Paracentral_Lobule_R<br>(25%) Precentral_R<br>(19%) OUTSIDE<br>(15%) Postcentral_R<br>(11%) Paracentral_Lobule_L | < 0.001 | < 0.001 | < 0.001 | 5,13 | **SM** |
| -38 | -2 | -14 | 320 | (36%) Rolandic_Oper_L<br>(32%) Insula_L<br>(14%) Postcentral_L<br>(13%) Temporal_Sup_L | < 0.001 | < 0.001 | < 0.001 | 4,99 | **Oper L** |

Functional regions are reported with corresponding MNI coordinates of peak activity within each cluster as well as cluster size. Additionally noted is the percentage of cluster's voxel in each anatomical region defined using the AAL3 atlas. Only regions with more than 5% of all cluster's voxel are noted. Results are thresholded at $p<0.001$ for voxel and FWE-corrected ($p<0.05$) for cluster size. Functional labels which are visible on the figures are marked in bold.

Table S3

Regions differing in BOLD activity between multisource vs no-conflict conditions.

| Coordinates MNI x | y | z | Cluster size | (% of cluster) AAL Region | Cluster pFWE | Cluster pFDR | Cluster p(uncor) | Peak T | Cluster label |
|---|---|---|---|---|---|---|---|---|---|
| **FS > 00** | | | | | | | | | |
| -28 | -60 | 50 | 15659 | (13%) OUTSIDE<br>(13%) Precentral_L<br>(10%) Parietal_Sup_L<br>(10%) Parietal_Inf_L<br>(7%) Postcentral_L<br>(7%) Supp_Motor_Area_L<br>(6%) Supp_Motor_Area_R<br>(6%) Frontal_Sup_2_R | < 0.001 | < 0.001 | < 0.001 | 12,3 | **pMFC<br>+ IPS<br>+ FEF<br>+ vPM<br>+ Oper** |
| 36 | -50 | -32 | 13306 | (11%) OUTSIDE<br>(8%) Cerebelum_6_R<br>(7%) Cerebelum_6_L<br>(7%) Cerebelum_8_R<br>(7%) Occipital_Inf_R<br>(6%) Cerebelum_Crus1_L<br>(6%) Occipital_Inf_L | < 0.001 | < 0.001 | < 0.001 | 11,33 | **LOC (R+L)** |
| 32 | 20 | 8 | 908 | (50%) Insula_R<br>(20%) OUTSIDE<br>(16%) Frontal_Inf_Oper_R<br>(7%) Frontal_Inf_Tri_R | < 0.001 | < 0.001 | < 0.001 | 11,17 | **aIns R<br>+ Oper** |
| 28 | -56 | 48 | 4092 | (29%) Parietal_Sup_R<br>(16%) Parietal_Inf_R<br>(15%) OUTSIDE<br>(10%) Precuneus_R<br>(10%) Postcentral_R<br>(6%) Occipital_Sup_R<br>(6%) Angular_R | < 0.001 | < 0.001 | < 0.001 | 10,14 | **IPS R<br>+ Prec R<br>+ Ang R** |
| -32 | 16 | 8 | 545 | (76%) Insula_L | < 0.001 | < 0.001 | < 0.001 | 9,18 | **aIns L** |

| | | | | | | | | | |
|---|---|---|---|---|---|---|---|---|---|
| | | | | (11%) OUTSIDE | | | | | |
| 34 | 54 | 26 | 1491 | (67%) Frontal_Mid_2_R<br>(16%) OUTSIDE<br>(11%) Frontal_Sup_2_R<br>(6%) Frontal_Inf_Tri_R | < 0.001 | < 0.001 | < 0.001 | 7,92 | **dlPFC R** |
| 14 | -12 | 10 | 619 | (28%) OUTSIDE<br>(25%) Caudate_R<br>(22%) Thal_VL_R<br>(6%) Thal_VPL_R | < 0.001 | < 0.001 | < 0.001 | 6,26 | Thal R<br>Caud R |
| 4 | -24 | -14 | 235 | (72%) OUTSIDE<br>(6%) Red_N_R<br>(6%) SN_pc_L | 0 | < 0.001 | < 0.001 | 6,16 | |
| -44 | 28 | 26 | 875 | (54%) Frontal_Mid_2_L<br>(37%) Frontal_Inf_Tri_L<br>(8%) OUTSIDE | < 0.001 | < 0.001 | < 0.001 | 6,07 | **dlPFC L** |
| -12 | -12 | 10 | 537 | (30%) Thal_VL_L<br>(24%) OUTSIDE<br>(12%) Caudate_L<br>(8%) Pallidum_L<br>(7%) Thal_VA_L<br>(6%) Thal_VPL_L | < 0.001 | < 0.001 | < 0.001 | 5,61 | Thal L<br>Caud L |
| **FS < 00** | | | | | | | | | |
| 18 | -90 | 22 | 48224 | (12%) OUTSIDE<br>(7%) Temporal_Mid_L<br>(5%) Temporal_Mid_R | < 0.001 | < 0.001 | < 0.001 | 9,81 | **aTmp<br>+Ling<br>+ Hip<br>+ Cun<br>+Prec<br>+vmPFC<br>+vlPFC<br>+dmPFC<br>+Ang<br>+Oper<br>+SM** |

| | | | | | | | | |
|---|---|---|---|---|---|---|---|---|
| 26 | -82 | -38 | 861 | (74%) Cerebelum_Crus2_R<br>(18%) Cerebelum_Crus1_R<br>(8%) OUTSIDE | < 0.001 | < 0.001 | < 0.001 | 6,61 |
| -16 | -86 | -38 | 211 | (81%) Cerebelum_Crus2_L<br>(17%) OUTSIDE | 0 | 0 | < 0.001 | 5,67 |

Functional regions are reported with corresponding MNI coordinates of peak activity within each cluster as well as cluster size. Additionally noted is the percentage of cluster's voxel in each anatomical region defined using the AAL3 atlas. Only regions with more than 5% of all cluster's voxel are noted. Results are thresholded at p<0.001 for voxel and FWE-corrected (p<0.05) for cluster size. Functional labels which are visible on the figures are marked in bold.

Table S4
Conjunction of flanker and Simon effects.

| Coordinates MNI x | y | z | Cluster size | (% of cluster) AAL Region | Cluster pFWE | Cluster pFDR | Cluster p(uncor) | Peak T | Cluster label |
|---|---|---|---|---|---|---|---|---|---|
| **(S0 > 00) ∩ (F0 > 00)** | | | | | | | | | |
| -28 | 4 | 60 | 4463 | (18%) Precentral_L<br>(15%) Frontal_Sup_2_R<br>(13%) Supp_Motor_Area_L<br>(13%) Frontal_Sup_2_L<br>(13%) Supp_Motor_Area_R<br>(12%) OUTSIDE | < 0.001 | < 0.001 | < 0.001 | 6,74 | **FEF +pMFC +vPM (l)** |
| -26 | -58 | 50 | 3189 | (31%) Parietal_Inf_L<br>(30%) Parietal_Sup_L<br>(12%) Postcentral_L<br>(10%) OUTSIDE<br>(8%) Precuneus_L<br>(6%) Occipital_Mid_L | < 0.001 | < 0.001 | < 0.001 | 6,51 | **IPS L** |
| 32 | -54 | 48 | 1778 | (34%) Parietal_Sup_R<br>(14%) OUTSIDE<br>(14%) Parietal_Inf_R<br>(9%) Precuneus_R<br>(7%) Occipital_Sup_R<br>(7%) Angular_R<br>(7%) Postcentral_R<br>(5%) SupraMarginal_R | < 0.001 | < 0.001 | < 0.001 | 4,63 | **IPS R** |
| -34 | -52 | -28 | 462 | (62%) Cerebelum_6_L<br>(34%) Cerebelum_Crus1_L | 0,03 | 0 | < 0.001 | 4,49 | |
| -38 | 38 | 28 | 466 | (56%) Frontal_Mid_2_L<br>(38%) Frontal_Inf_Tri_L<br>(5%) OUTSIDE | 0,03 | 0 | < 0.001 | 4,13 | **dlPFC L** |
| 36 | 54 | 32 | 522 | (74%) Frontal_Mid_2_R<br>(14%) OUTSIDE<br>(12%) Frontal_Sup_2_R | 0,02 | 0 | < 0.001 | 3,7 | **dlPFC R** |

| | | | | | | | | | |
|---|---|---|---|---|---|---|---|---|---|
| **(S0 < 00) ∩ (F0 < 00)** | | | | | | | | | |
| -2 | 62 | -8 | 2792 | (34%) Frontal_Sup_Medial_L<br>(25%) Frontal_Sup_2_L<br>(12%) Frontal_Med_Orb_L<br>(10%) Frontal_Sup_Medial_R<br>(5%) OUTSIDE<br>(5%) Frontal_Med_Orb_R | < 0.001 | < 0.001 | < 0.001 | 5,83 | **vlPFC<br>+ vmPFC<br>+ dmPFC** |
| -60 | -64 | 32 | 858 | (64%) Angular_L<br>(22%) OUTSIDE | 0 | < 0.001 | < 0.001 | 5,06 | **Ang L** |
| 54 | -70 | 38 | 536 | (48%) Angular_R<br>(29%) Temporal_Mid_R<br>(8%) OUTSIDE<br>(7%) Occipital_Mid_R<br>(6%) Temporal_Sup_R | 0,02 | 0 | < 0.001 | 4,31 | **Ang R** |
| 58 | 0 | -28 | 940 | (33%) Temporal_Mid_R<br>(25%) Temporal_Pole_Mid_R<br>(21%) Temporal_Pole_Sup_R<br>(6%) ParaHippocampal_R<br>(5%) OFCpost_R<br>(5%) Temporal_Inf_R | < 0.001 | < 0.001 | < 0.001 | 4,24 | **aTmpR** |
| -64 | -28 | -14 | 873 | (52%) Temporal_Mid_L<br>(38%) OUTSIDE<br>(9%) Temporal_Inf_L | < 0.001 | < 0.001 | < 0.001 | 4,14 | **aTmp L** |
| -38 | 20 | -26 | 1051 | (29%) Frontal_Inf_Tri_L<br>(18%) Frontal_Inf_Orb_2_L<br>(16%) OFCpost_L<br>(15%) Temporal_Pole_Sup_L<br>(11%) OUTSIDE<br>(10%) Hippocampus_R | < 0.001 | < 0.001 | < 0.001 | 4,12 | vlPFC |
| 4 | -54 | 18 | 886 | (23%) Precuneus_L<br>(22%) Precuneus_R<br>(15%) Cingulate_Mid_L<br>(11%) Cingulate_Post_L | < 0.001 | < 0.001 | < 0.001 | 3,89 | **Prec** |

| | |
|---|---|
| | (9%) Cingulate_Mid_R |
| | (6%) Calcarine_L |

Functional regions are reported with corresponding MNI coordinates of peak activity within each cluster as well as cluster size. Additionally noted is the percentage of cluster's voxel in each anatomical region defined using the AAL3 atlas. Only regions with more than 5% of all cluster's voxel are noted. Results are thresholded at p<0.001 for voxel and FWE-corrected (p<0.05) for cluster size. Functional labels which are visible on the figures are marked in bold.

Table S5

Difference between flanker and Simon conditions.

| Coordinates MNI x | y | z | Cluster size | (% of cluster) AAL Region | Cluster pFWE | Cluster pFDR | Cluster p(uncor) | Peak T | Cluster label |
|---|---|---|---|---|---|---|---|---|---|
| **F0 > S0** | | | | | | | | | |
| 36 | -90 | -2 | 4758 | (16%) Occipital_Inf_R<br>(12%) Occipital_Mid_R<br>(11%) OUTSIDE<br>(9%) Parietal_Sup_R<br>(9%) Cerebelum_Crus1_R<br>(8%) Cerebelum_6_R<br>(6%) Parietal_Inf_R | < 0.001 | < 0.001 | < 0.001 | 10,39 | **LOC R**<br>**LOC R** |
| -32 | -92 | -4 | 2440 | (23%) Cerebelum_Crus1_L<br>(22%) Occipital_Inf_L<br>(15%) Occipital_Mid_L<br>(14%) OUTSIDE<br>(10%) Cerebelum_6_L<br>(5%) Lingual_L | < 0.001 | < 0.001 | < 0.001 | 8,33 | **LOC L** |
| -24 | -64 | 44 | 2003 | (38%) Parietal_Inf_L<br>(33%) Parietal_Sup_L<br>(11%) Occipital_Mid_L<br>(9%) OUTSIDE | < 0.001 | < 0.001 | < 0.001 | 7,45 | **IPS L** |
| -44 | 0 | 34 | 1087 | (70%) Precentral_L<br>(9%) Frontal_Mid_2_L<br>(8%) Frontal_Inf_Oper_L<br>(6%) Frontal_Sup_2_L | < 0.001 | < 0.001 | < 0.001 | 7,35 | **vPM L** |
| -4 | 14 | 50 | 884 | (39%) Supp_Motor_Area_L<br>(21%) Supp_Motor_Area_R<br>(18%) Cingulate_Mid_R<br>(8%) Cingulate_Mid_L<br>(8%) Frontal_Sup_Medial_L | < 0.001 | < 0.001 | < 0.001 | 7,08 | **pMFC** |
| -14 | -16 | 16 | 203 | (38%) Thal_VL_L<br>(16%) Thal_VA_L<br>(11%) OUTSIDE | 0 | 0 | < 0.001 | 6,12 | Thal L |

| | | | | | | | | | |
|---|---|---|---|---|---|---|---|---|---|
| | | | | (9%) Thal_VPL_L<br>(6%) Thal_PuM_L<br>(6%) Thal_MDl_L | | | | | |
| 28 | -70 | -50 | 134 | (84%) Cerebelum_8_R<br>(16%) Cerebelum_7b_R | 0,03 | 0 | 0 | 5,82 | |
| 44 | 34 | 30 | 203 | (68%) Frontal_Mid_2_R<br>(23%) Frontal_Inf_Tri_R<br>(8%) OUTSIDE | 0 | 0 | < 0.001 | 5,52 | **dlPFC** |
| -30 | -70 | -50 | 125 | (42%) Cerebelum_7b_L<br>(37%) Cerebelum_8_L<br>(21%) Cerebelum_Crus2_L | 0,04 | 0 | 0 | 5,44 | |
| 40 | 20 | -4 | 138 | (88%) Insula_R<br>(5%) Frontal_Inf_Orb_2_R | 0,02 | 0 | 0 | 5,15 | **aIns R** |
| 26 | -2 | 50 | 166 | (59%) Frontal_Sup_2_R<br>(23%) Frontal_Mid_2_R<br>(11%) Precentral_R<br>(6%) OUTSIDE | 0,01 | 0 | < 0.001 | 5,14 | **FEF R** |
| 46 | 6 | 34 | 207 | (58%) Precentral_R<br>(39%) Frontal_Inf_Oper_R | 0 | 0 | < 0.001 | 4,95 | **vPM R** |
| **F0 < S0** | | | | | | | | | |
| -12 | -80 | -10 | 829 | (76%) Lingual_L<br>(14%) Fusiform_L<br>(7%) Cerebelum_6_L | < 0.001 | < 0.001 | < 0.001 | 7,55 | **Ling L** |
| 18 | -76 | -12 | 874 | (81%) Lingual_R<br>(15%) Fusiform_R | < 0.001 | < 0.001 | < 0.001 | 7,35 | **Ling R** |
| -12 | -92 | 16 | 1673 | (27%) Cuneus_L<br>(20%) Cuneus_R<br>(20%) Occipital_Sup_L<br>(11%) Calcarine_L<br>(9%) Calcarine_R<br>(6%) Occipital_Sup_R | < 0.001 | < 0.001 | < 0.001 | 6,59 | **Cun<br>+ Cal** |
| 54 | -60 | 6 | 1408 | (41%) Temporal_Mid_R | < 0.001 | < 0.001 | < 0.001 | 6,4 | **Ang R** |

| | | | | | | | | | |
|---|---|---|---|---|---|---|---|---|---|
| | | | | (37%) Angular_R<br>(13%) Occipital_Mid_R<br>(5%) OUTSIDE | | | | | **+ pTmp R** |
| -60 | -68 | 18 | 742 | (51%) OUTSIDE<br>(24%) Angular_L<br>(17%) Occipital_Mid_L | < 0.001 | < 0.001 | < 0.001 | 5,81 | **Ang L** |
| -50 | -8 | -16 | 182 | (67%) Temporal_Mid_L<br>(20%) Temporal_Sup_L<br>(8%) Temporal_Pole_Sup_L | 0,01 | 0 | < 0.001 | 5,77 | **aTmp L** |
| 62 | -26 | 28 | 782 | (46%) SupraMarginal_R<br>(35%) Rolandic_Oper_R<br>(5%) Postcentral_R | < 0.001 | < 0.001 | < 0.001 | 5,57 | **SMG R<br>+ Oper** |
| -60 | -30 | 24 | 192 | (72%) SupraMarginal_L<br>(23%) Temporal_Sup_L | 0 | 0 | < 0.001 | 5,55 | **SMG L** |
| -40 | 0 | 8 | 418 | (33%) Postcentral_L<br>(23%) Insula_L<br>(22%) Rolandic_Oper_L<br>(9%) Precentral_L<br>(6%) Frontal_Inf_Oper_L<br>(6%) OUTSIDE | < 0.001 | < 0.001 | < 0.001 | 5,31 | SM<br>+ Oper |
| 8 | -32 | 46 | 418 | (50%) Cingulate_Mid_R<br>(33%) Cingulate_Mid_L<br>(6%) Paracentral_Lobule_R | < 0.001 | < 0.001 | < 0.001 | 5,27 | **MCC (R+L)** |
| 0 | -32 | 66 | 134 | (55%) Paracentral_Lobule_R<br>(27%) Paracentral_Lobule_L<br>(18%) OUTSIDE | 0,03 | 0,01 | 0 | 5,23 | SM |
| 52 | -8 | -16 | 234 | (59%) Temporal_Mid_R<br>(28%) Temporal_Sup_R<br>(7%) Temporal_Pole_Sup_R | 0 | < 0.001 | < 0.001 | 5,11 | **aTmp R** |
| 22 | 26 | 60 | 325 | (93%) Frontal_Sup_2_R<br>(7%) Frontal_Mid_2_R | < 0.001 | < 0.001 | < 0.001 | 4,82 | **dmPFC R** |
| -24 | 32 | 46 | 194 | (66%) Frontal_Mid_2_L | 0 | 0 | < 0.001 | 4,63 | **dmPFC L** |

| | | | | | | | | | |
|---|---|---|---|---|---|---|---|---|---|
| | | | | (34%) Frontal_Sup_2_L | | | | | |
| 32 | -36 | -10 | 121 | (59%) ParaHippocampal_R<br>(29%) Fusiform_R<br>(11%) Hippocampus_R | 0,04 | 0,01 | 0 | 4,61 | Hip R |
| 28 | -40 | 64 | 122 | (86%) Postcentral_R<br>(10%) Parietal_Sup_R | 0,04 | 0,01 | 0 | 4,34 | SM |

Functional regions are reported with corresponding MNI coordinates of peak activity within each cluster as well as cluster size. Additionally noted is the percentage of cluster's voxel in each anatomical region defined using the AAL3 atlas. Only regions with more than 5% of all cluster's voxel are noted. Results are thresholded at p<0.001 for voxel and FWE-corrected (p<0.05) for cluster size. Functional labels which are visible on the figures are marked in bold.

Table S6

Difference between flanker and Simon effects masked by their conjunction map.

| Coordinates MNI x | y | z | Cluster size | (% of cluster) AAL Region | Cluster pFWE | Cluster pFDR | Cluster p(uncor) | Peak T | Cluster label |
|---|---|---|---|---|---|---|---|---|---|
| **F0 > S0 (mask with S0 ∩ F0)** | | | | | | | | | |
| 36 | -90 | -2 | 2215 | (30%) Occipital_Inf_R<br>(24%) Occipital_Mid_R<br>(14%) OUTSIDE<br>(8%) Lingual_R<br>(7%) Parietal_Sup_R<br>(5%) Occipital_Sup_R | < 0.001 | < 0.001 | < 0.001 | 10,39 | **LOC R + IPS R** |
| -32 | -92 | -4 | 1899 | (23%) Cerebelum_Crus1_L<br>(21%) Occipital_Inf_L<br>(17%) OUTSIDE<br>(15%) Occipital_Mid_L<br>(7%) Cerebelum_6_L<br>(6%) Lingual_L | < 0.001 | < 0.001 | < 0.001 | 8,33 | **LOC L + Ling L** |
| -4 | -76 | -30 | 1360 | (31%) Cerebelum_Crus1_R<br>(24%) Cerebelum_6_R<br>(10%) Cerebelum_Crus2_L<br>(9%) Vermis_7<br>(5%) Cerebelum_Crus1_L<br>(5%) Cerebelum_Crus2_R | < 0.001 | < 0.001 | < 0.001 | 6,9 | |
| -28 | -70 | 54 | 227 | (44%) Parietal_Sup_L<br>(19%) Precuneus_L<br>(12%) Parietal_Inf_L<br>(11%) Occipital_Sup_L<br>(7%) OUTSIDE<br>(6%) Occipital_Mid_L | 0 | 0 | < 0.001 | 6,15 | **IPS L** |
| -14 | -16 | 16 | 203 | (38%) Thal_VL_L<br>(16%) Thal_VA_L<br>(11%) OUTSIDE<br>(9%) Thal_VPL_L | 0 | 0 | < 0.001 | 6,12 | **Thal L** |

| | | | | | | | | | |
|---|---|---|---|---|---|---|---|---|---|
| | | | | (6%) Thal_PuM_L<br>(6%) Thal_MDl_L | | | | | |
| 8 | 24 | 34 | 355 | (25%) Supp_Motor_Area_L<br>(19%) Cingulate_Mid_R<br>(18%) Cingulate_Mid_L<br>(17%) Frontal_Sup_Medial_L<br>(13%) Supp_Motor_Area_R<br>(6%) ACC_sup_L | < 0.001 | 0 | < 0.001 | 5,56 | **pMFC** |
| -38 | 6 | 32 | 406 | (67%) Precentral_L<br>(19%) Frontal_Inf_Oper_L<br>(5%) Frontal_Mid_2_L | < 0.001 | 0 | < 0.001 | 5,53 | **FEF** |
| 44 | 34 | 30 | 186 | (66%) Frontal_Mid_2_R<br>(25%) Frontal_Inf_Tri_R<br>(9%) OUTSIDE | 0,01 | 0 | < 0.001 | 5,52 | **dlPFC** |
| -30 | -70 | -50 | 125 | (42%) Cerebelum_7b_L<br>(37%) Cerebelum_8_L<br>(21%) Cerebelum_Crus2_L | 0,04 | 0 | 0 | 5,44 | |
| **F0 < S0 (mask with S0 ∩ F0)** | | | | | | | | | |
| -12 | -80 | -10 | 811 | (76%) Lingual_L<br>(14%) Fusiform_L<br>LOC L + Ling L | < 0.001 | < 0.001 | < 0.001 | 7,55 | **Ling L** |
| 18 | -76 | -12 | 862 | (80%) Lingual_R<br>(16%) Fusiform_R | < 0.001 | < 0.001 | < 0.001 | 7,35 | **Ling R** |
| -12 | -92 | 16 | 1480 | (30%) Cuneus_L<br>(22%) Occipital_Sup_L<br>(18%) Cuneus_R<br>(11%) Calcarine_L<br>(5%) Calcarine_R | < 0.001 | < 0.001 | < 0.001 | 6,59 | **Cun<br>+ Cal** |
| 54 | -60 | 6 | 1149 | (47%) Temporal_Mid_R<br>(32%) Angular_R<br>(14%) Occipital_Mid_R<br>(5%) OUTSIDE | < 0.001 | < 0.001 | < 0.001 | 6,4 | **Ang R** |

| | | | | | | | | | |
|---|---|---|---|---|---|---|---|---|---|
| -60 | -68 | 18 | 565 | (60%) OUTSIDE<br>(19%) Occipital_Mid_L<br>(9%) Angular_L<br>(6%) Temporal_Mid_L | < 0.001 | < 0.001 | < 0.001 | 5,81 | **Ang L** |
| -50 | -8 | -16 | 174 | (66%) Temporal_Mid_L<br>(21%) Temporal_Sup_L<br>(9%) Temporal_Pole_Sup_L<br>(5%) Insula_L | 0,01 | 0,01 | < 0.001 | 5,77 | **aTmp L** |
| 62 | -26 | 28 | 782 | (46%) SupraMarginal_R<br>(35%) Rolandic_Oper_R<br>(5%) Postcentral_R | < 0.001 | < 0.001 | < 0.001 | 5,57 | **SMG R** |
| -60 | -30 | 24 | 192 | (72%) SupraMarginal_L<br>(23%) Temporal_Sup_L | 0 | 0 | < 0.001 | 5,55 | **SMG L** |
| -40 | 0 | 8 | 418 | (33%) Postcentral_L<br>(23%) Insula_L<br>(22%) Rolandic_Oper_L<br>(9%) Precentral_L<br>(6%) Frontal_Inf_Oper_L<br>(6%) OUTSIDE | < 0.001 | < 0.001 | < 0.001 | 5,31 | SM L<br>+ Oper L<br>+ Ins L |
| 8 | -32 | 46 | 416 | (50%) Cingulate_Mid_R<br>(33%) Cingulate_Mid_L<br>(6%) Paracentral_Lobule_R | < 0.001 | < 0.001 | < 0.001 | 5,27 | **MCC** |
| 0 | -32 | 66 | 134 | (55%) Paracentral_Lobule_R<br>(27%) Paracentral_Lobule_L<br>(18%) OUTSIDE | 0,03 | 0,01 | 0 | 5,23 | **MCC** |
| 52 | -8 | -16 | 182 | (52%) Temporal_Mid_R<br>(32%) Temporal_Sup_R<br>(9%) Temporal_Pole_Sup_R<br>(5%) Temporal_Pole_Mid_R | 0,01 | 0 | < 0.001 | 5,11 | **Tmp R** |
| 22 | 26 | 60 | 314 | (92%) Frontal_Sup_2_R<br>(7%) Frontal_Mid_2_R | < 0.001 | < 0.001 | < 0.001 | 4,82 | **dmPFC R** |
| -24 | 32 | 46 | 189 | (67%) Frontal_Mid_2_L | 0 | 0 | < 0.001 | 4,63 | **dmPFC L** |

| | | | | | | | | | |
|---|---|---|---|---|---|---|---|---|---|
| | | | | (33%) Frontal_Sup_2_L | | | | | |
| 28 | -40 | 64 | 122 | (86%) Postcentral_R | 0,04 | 0,01 | 0 | 4,34 | SM |
| | | | | (10%) Parietal_Sup_R | | | | | |

Functional regions are reported with corresponding MNI coordinates of peak activity within each cluster as well as cluster size. Additionally noted is the percentage of cluster's voxel in each anatomical region defined using the AAL3 atlas. Only regions with more than 5% of all cluster's voxel are noted. Results are thresholded at p<0.001 for voxel and FWE-corrected (p<0.05) for cluster size. Functional labels which are visible on the figures are marked in bold.

Table S7

Regions exceeding the predicted time-on-task effects in Simon condition.

| Coordinates MNI x | y | z | Cluster size | (% of cluster) AAL Region | Cluster pFWE | Cluster pFDR | Cluster p(uncor) | Peak T | Cluster label |
|---|---|---|---|---|---|---|---|---|---|
| **S0 > rtS0** | | | | | | | | | |
| -24 | -52 | 42 | 126 | (43%) Parietal_Inf_L<br>(35%) Parietal_Sup_L<br>(22%) OUTSIDE | 0,05 | 0,07 | 0 | 4,76 | **IPS L** |
| **S0 < rtS0** | | | | | | | | | |
| - | - | - | - | - | - | - | - | - | |

Functional regions are reported with corresponding MNI coordinates of peak activity within each cluster as well as cluster size. Additionally noted is the percentage of cluster's voxel in each anatomical region defined using the AAL3 atlas. Only regions with more than 5% of all cluster's voxel are noted. Results are thresholded at $p<0.001$ for voxel and FWE-corrected ($p<0.05$) for cluster size. Functional labels which are visible on the figures are marked in bold.

Table S8

Regions exceeding the predicted time-on-task effects in flanker condition.

| Coordinates MNI x | y | z | Cluster size | (% of cluster) AAL Region | Cluster pFWE | Cluster pFDR | Cluster p(uncor) | Peak T | Cluster label |
|---|---|---|---|---|---|---|---|---|---|
| **F0 > rtF0** | | | | | | | | | |
| -36 | -50 | -2 | 466 | (90%) OUTSIDE | < 0.001 | < 0.001 | < 0.001 | 6,33 | wm L |
| 24 | -32 | 10 | 971 | (86%) OUTSIDE | < 0.001 | < 0.001 | < 0.001 | 6,21 | wm R<br>**IPS** |
| -22 | -70 | -34 | 1759 | (46%) Cerebelum_Crus1_L<br>(20%) Cerebelum_Crus2_L<br>(12%) Cerebelum_8_L<br>(8%) Cerebelum_6_L<br>(6%) OUTSIDE | < 0.001 | < 0.001 | < 0.001 | 5,93 | |
| -22 | 12 | -2 | 937 | (61%) OUTSIDE<br>(23%) Putamen_L<br>(11%) Caudate_L | < 0.001 | < 0.001 | < 0.001 | 5,91 | Thal L<br>Caud L |
| 20 | 0 | 22 | 2786 | (57%) OUTSIDE<br>(16%) Caudate_R<br>(10%) Putamen_R | < 0.001 | < 0.001 | < 0.001 | 5,88 | **vlPFC**<br>**+** aIns R<br>Thal R<br>Caud R |
| 6 | 14 | 60 | 512 | (72%) Supp_Motor_Area_R<br>(22%) Supp_Motor_Area_L | < 0.001 | < 0.001 | < 0.001 | 5,62 | **pMFC** |
| -22 | -48 | 36 | 500 | (84%) OUTSIDE<br>(11%) Parietal_Sup_L | < 0.001 | < 0.001 | < 0.001 | 5,43 | wm L<br>**IPS** |
| 30 | -56 | -38 | 310 | (44%) Cerebelum_Crus1_R<br>(23%) Cerebelum_6_R<br>(20%) Cerebelum_8_R<br>(14%) OUTSIDE | < 0.001 | < 0.001 | < 0.001 | 5,16 | |
| 26 | 58 | 32 | 135 | (71%) Frontal_Sup_2_R<br>(15%) OUTSIDE<br>(14%) Frontal_Mid_2_R | 0,02 | 0,01 | 0 | 4,96 | **dlPFC R** |
| 38 | -90 | 0 | 224 | (27%) OUTSIDE<br>(26%) Occipital_Inf_R | 0 | 0 | < 0.001 | 4,37 | **LOC R** |

| | | | | | | | | | |
|---|---|---|---|---|---|---|---|---|---|
| | | | | (26%) Occipital_Mid_R<br>(13%) Calcarine_R<br>(6%) Fusiform_R | | | | | |
| -26 | -2 | 26 | 212 | (93%) OUTSIDE | 0 | 0 | < 0.001 | 4,34 | |
| **F0 < rtF0** | | | | | | | | | |
| 16 | -58 | 14 | 1213 | (28%) Calcarine_R<br>(22%) Lingual_R<br>(21%) Calcarine_L<br>(11%) Lingual_L<br>(8%) Cuneus_L | < 0.001 | < 0.001 | < 0.001 | 5,71 | **Cal<br>+ Ling<br>+ Cun L** |
| 68 | -8 | 8 | 168 | (49%) Rolandic_Oper_R<br>(39%) Temporal_Sup_R<br>(7%) Heschl_R | 0,01 | 0 | < 0.001 | 5,58 | Oper |
| 26 | -42 | -16 | 333 | (83%) Fusiform_R<br>(11%) Cerebelum_6_R<br>(6%) Cerebelum_4_5_R | < 0.001 | < 0.001 | < 0.001 | 5,37 | **Fus R** |
| -28 | -88 | 36 | 148 | (70%) Occipital_Mid_L<br>(14%) OUTSIDE<br>(10%) Parietal_Inf_L<br>(7%) Occipital_Sup_L | 0,02 | 0 | < 0.001 | 5,06 | **LOC L** |
| -44 | -52 | -22 | 238 | (82%) Fusiform_L<br>(6%) Cerebelum_6_L<br>(5%) Temporal_Inf_L | 0 | 0 | < 0.001 | 5,03 | **Fus L** |
| 38 | -22 | 66 | 166 | (100%) Precentral_R | 0,01 | 0 | < 0.001 | 4,92 | **SM R** |
| -60 | -4 | 18 | 143 | (87%) Postcentral_L<br>(13%) Precentral_L | 0,02 | 0,01 | 0 | 4,91 | **SM L** |
| 58 | -14 | 46 | 125 | (77%) Postcentral_R<br>(22%) Precentral_R | 0,04 | 0,01 | 0 | 4,71 | **SM R** |

Functional regions are reported with corresponding MNI coordinates of peak activity within each cluster as well as cluster size. Additionally noted is the percentage of cluster's voxel in each anatomical region defined using the AAL3 atlas. Only regions with more than 5% of all cluster's voxel are noted. Results are thresholded at p<0.001 for voxel and FWE-corrected (p<0.05) for cluster size. Functional labels which are visible on the figures are marked in bold.

Table S9

Regions exceeding the predicted time-on-task effects in multi-source condition.

| Coordinates MNI x | y | z | Cluster size | (% of cluster) AAL Region | Cluster pFWE | Cluster pFDR | Cluster p(uncor) | Peak T | Cluster label |
|---|---|---|---|---|---|---|---|---|---|
| **FS > rtFS** | | | | | | | | | |
| -36 | -50 | -2 | 336 | (92%) OUTSIDE | < 0.001 | < 0.001 | < 0.001 | 6,26 | wm L |
| 20 | -20 | 26 | 2427 | (57%) OUTSIDE<br>(16%) Caudate_R<br>(12%) Putamen_R | < 0.001 | < 0.001 | < 0.001 | 6,02 | wm<br>Caud<br>Put<br>+ aIns R |
| 10 | 18 | 68 | 481 | (79%) Supp_Motor_Area_R<br>(10%) Frontal_Sup_Medial_R<br>(5%) OUTSIDE | < 0.001 | < 0.001 | < 0.001 | 5,9 | **pMFC R** |
| -12 | -44 | -42 | 1033 | (43%) Cerebelum_Crus1_L<br>(20%) Cerebelum_8_L<br>(13%) Cerebelum_Crus2_L<br>(10%) Cerebelum_6_L<br>(6%) OUTSIDE<br>(5%) Cerebelum_9_L | < 0.001 | < 0.001 | < 0.001 | 5,6 | |
| -22 | 12 | -2 | 401 | (46%) OUTSIDE<br>(35%) Putamen_L<br>(16%) Caudate_L | < 0.001 | < 0.001 | < 0.001 | 5,53 | wm<br>Caud<br>Put |
| 38 | -44 | 4 | 1087 | (91%) OUTSIDE | < 0.001 | < 0.001 | < 0.001 | 5,52 | wm |
| 30 | -56 | -38 | 134 | (58%) Cerebelum_8_R<br>(25%) OUTSIDE<br>(9%) Cerebelum_6_R<br>(8%) Cerebelum_Crus1_R | 0,03 | 0,01 | 0 | 5,2 | |
| -16 | 10 | 20 | 226 | (61%) OUTSIDE<br>(38%) Caudate_L | 0 | 0 | < 0.001 | 5,01 | |
| -28 | -18 | 32 | 622 | (97%) OUTSIDE | < 0.001 | < 0.001 | < 0.001 | 4,97 | |
| -22 | -50 | 38 | 258 | (94%) OUTSIDE<br>(6%) Parietal_Sup_L<br>(24%) Cerebelum_6_R | 0 | < 0.001 | < 0.001 | 4,8 | wm |

| | | | | | | | | | |
|---|---|---|---|---|---|---|---|---|---|
| **FS < rtFS** | | | | | | | | | |
| 40 | -22 | 64 | 350 | (81%) Precentral_R<br>(14%) Postcentral_R | < 0.001 | < 0.001 | < 0.001 | 5,77 | **SM** |
| 68 | -8 | 6 | 273 | (42%) Temporal_Sup_R<br>(34%) Rolandic_Oper_R<br>(16%) Temporal_Pole_Sup_R<br>(7%) Heschl_R | 0 | < 0.001 | < 0.001 | 5,61 | Oper |
| 14 | -54 | -2 | 1823 | (16%) Lingual_R<br>(16%) Calcarine_L<br>(15%) Calcarine_R<br>(11%) Cuneus_L<br>(10%) Lingual_L<br>(9%) Occipital_Sup_L<br>(7%) Cuneus_R | < 0.001 | < 0.001 | < 0.001 | 5,57 | **Cal<br>+ Ling<br>+ Cun** |
| 34 | -62 | -18 | 227 | (87%) Fusiform_R<br>(10%) Cerebelum_6_R | 0 | 0 | < 0.001 | 4,98 | **Fus R** |
| -44 | -52 | -22 | 455 | (44%) Fusiform_L<br>(35%) Temporal_Inf_L<br>(9%) Occipital_Inf_L<br>(6%) Temporal_Mid_L | < 0.001 | < 0.001 | < 0.001 | 4,71 | **Fus L** |

Functional regions are reported with corresponding MNI coordinates of peak activity within each cluster as well as cluster size. Additionally noted is the percentage of cluster's voxel in each anatomical region defined using the AAL3 atlas. Only regions with more than 5% of all cluster's voxel are noted. Results are thresholded at p<0.001 for voxel and FWE-corrected (p<0.05) for cluster size. Functional labels which are visible on the figures are marked in bold.

Conjunction of flanker and Simon activity exceeding time on task effects.

| Coordinates MNI x | y | z | Cluster size | (% of cluster) AAL Region | Cluster pFWE | Cluster pFDR | Cluster p(uncor) | Peak T | Cluster label |
|---|---|---|---|---|---|---|---|---|---|
| **(S0 > rtS0) ∩ (F0 > rtF0)** | | | | | | | | | |
| 24 | 10 | 4 | 1326 | (31%) Putamen_R<br>(27%) OUTSIDE<br>(20%) Caudate_R<br>(10%) Insula_R<br>(8%) Frontal_Inf_Oper_R | < 0.001 | < 0.001 | < 0.001 | 4,77 | Caud<br>Put<br>wm<br>**aIns R**<br>+ Oper<br>**+vlPFC** |
| 12 | 14 | 66 | 772 | (59%) Supp_Motor_Area_R<br>(14%) Supp_Motor_Area_L<br>(9%) Cingulate_Mid_R<br>(8%) OUTSIDE<br>(6%) ACC_sup_R | 0 | 0 | < 0.001 | 4,49 | **pMFC** |
| -30 | -56 | -46 | 681 | (37%) Cerebelum_Crus1_L<br>(27%) Cerebelum_8_L<br>(14%) Cerebelum_Crus2_L<br>(13%) Cerebelum_6_L<br>(8%) OUTSIDE | 0 | 0 | < 0.001 | 4,14 | |
| 28 | -58 | -36 | 420 | (46%) Cerebelum_8_R<br>(32%) OUTSIDE<br>(14%) Cerebelum_6_R<br>(8%) Cerebelum_Crus1_R | 0,05 | 0,01 | < 0.001 | 3,82 | |
| **(S0 < rtS0) ∩ (F0 < rtF0)** | | | | | | | | | |
| 16 | -58 | 14 | 629 | (28%) Calcarine_L<br>(21%) Calcarine_R<br>(19%) Lingual_L<br>(13%) Lingual_R<br>(9%) Precuneus_R | 0,01 | 0 | < 0.001 | 4,26 | **Cal**<br>+ Ling<br>+ Precun R |

Functional regions are reported with corresponding MNI coordinates of peak activity within each cluster as well as cluster size. Additionally noted is the percentage of cluster's voxel in each anatomical region defined using the AAL3 atlas. Only regions with more than 5% of all cluster's voxel are noted. Results are thresholded at p<0.001 for voxel and FWE-corrected (p<0.05) for cluster size. Functional labels which are visible on the figures are marked in bold.

Table S11

Difference between flanker and Simon-related activity exceeding time-on-task effect.

| Coordinates MNI x | y | z | Cluster size | (% of cluster) AAL Region | Cluster pFWE | Cluster pFDR | Cluster p(uncor) | Peak T | Cluster label |
|---|---|---|---|---|---|---|---|---|---|
| **(F0 > rtF0) > (S0 > rtS0)** | | | | | | | | | |
| 20 | 30 | -2 | 4562 | (81%) OUTSIDE<br>(6%) Caudate_R | < 0.001 | < 0.001 | < 0.001 | 7,35 | Caud<br>wm<br>**IPS** |
| 42 | 24 | -10 | 359 | (67%) Frontal_Inf_Orb_2_R<br>(19%) Insula_R<br>(6%) Frontal_Inf_Tri_R | < 0.001 | < 0.001 | < 0.001 | 6,77 | **vlPFC**<br>+ aIns |
| -32 | -70 | -28 | 901 | (54%) Cerebelum_Crus1_L<br>(29%) Cerebelum_Crus2_L<br>(7%) Cerebelum_6_L<br>(6%) Cerebelum_8_L | < 0.001 | < 0.001 | < 0.001 | 5,2 | |
| -22 | -50 | 12 | 639 | (84%) OUTSIDE<br>(6%) Occipital_Mid_L<br>(5%) Occipital_Inf_L | < 0.001 | < 0.001 | < 0.001 | 5,16 | LOC L |
| 40 | -66 | -30 | 169 | (79%) Cerebelum_Crus1_R<br>(21%) Cerebelum_6_R | 0,01 | 0 | < 0.001 | 5,13 | |
| -6 | -72 | -26 | 364 | (30%) Cerebelum_Crus2_L<br>(20%) Cerebelum_Crus1_L<br>(18%) OUTSIDE<br>(7%) Cerebelum_6_L<br>(6%) Vermis_8 | < 0.001 | < 0.001 | < 0.001 | 4,98 | |
| 38 | -90 | 0 | 350 | (34%) Occipital_Mid_R<br>(25%) Occipital_Inf_R<br>(24%) Calcarine_R<br>(13%) OUTSIDE | < 0.001 | < 0.001 | < 0.001 | 4,73 | **LOC R**<br>+ Cal R |
| 12 | 56 | 24 | 200 | (56%) Frontal_Sup_Medial_R<br>(43%) Frontal_Sup_2_R | 0 | 0 | < 0.001 | 4,28 | **dmPFC** |
| **(F0 > rtF0) < (S0 > rtS0)** | | | | | | | | | |
| 24 | -42 | -16 | 240 | (80%) Fusiform_R<br>(13%) Cerebelum_4_5_R | 0 | < 0.001 | < 0.001 | 5,63 | **Fus R** |

| | | | | | | | | | |
|---|---|---|---|---|---|---|---|---|---|
| 50 | -60 | 4 | 212 | (97%) Temporal_Mid_R | 0 | 0 | < 0.001 | 5,4 | **pTmp** |
| -34 | -50 | -22 | 335 | (82%) Fusiform_L<br>(12%) Cerebelum_6_L<br>(5%) Temporal_Inf_L | < 0.001 | < 0.001 | < 0.001 | 5,24 | **Fus L** |
| 28 | -60 | -16 | 200 | (81%) Fusiform_R<br>(14%) Cerebelum_6_R | 0 | 0 | < 0.001 | 5,12 | **Fus R** |
| -12 | -66 | 4 | 1269 | (28%) Calcarine_R<br>(26%) Lingual_R<br>(22%) Calcarine_L<br>(13%) Cuneus_L<br>(9%) Lingual_L | < 0.001 | < 0.001 | < 0.001 | 5,05 | **Cal**<br>+ Ling<br>**+ Cun** |
| 40 | -78 | 30 | 110 | (88%) Occipital_Mid_R<br>(12%) Angular_R | 0,05 | 0,01 | 0 | 4,91 | **Ang R** |
| -8 | -58 | 64 | 174 | (80%) Precuneus_L<br>(19%) Parietal_Sup_L | 0,01 | 0 | < 0.001 | 4,78 | **Prec L** |
| -58 | -8 | 38 | 152 | (64%) Postcentral_L<br>(36%) Precentral_L | 0,01 | 0 | < 0.001 | 4,62 | **SM L** |
| -20 | -82 | 50 | 168 | (32%) Occipital_Mid_L<br>(23%) Parietal_Sup_L<br>(19%) Occipital_Sup_L<br>(13%) OUTSIDE<br>(11%) Precuneus_L | 0,01 | 0 | < 0.001 | 4,56 | **Prec L** |
| 54 | -20 | 48 | 300 | (55%) Postcentral_R<br>(40%) SupraMarginal_R | < 0.001 | < 0.001 | < 0.001 | 4,47 | **SMG R**<br>**+ SM** |
| 40 | -12 | 60 | 122 | (98%) Precentral_R | 0,03 | 0,01 | 0 | 4,41 | **SM** |
| -56 | -24 | 16 | 133 | (47%) SupraMarginal_L<br>(32%) Temporal_Sup_L<br>(16%) Postcentral_L<br>(5%) Rolandic_Oper_L | 0,02 | 0 | 0 | 4,34 | **SMG L** |

Functional regions are reported with corresponding MNI coordinates of peak activity within each cluster as well as cluster size. Additionally noted is the percentage of cluster's voxel in each anatomical region defined using the AAL3 atlas. Only regions with more than 5% of all cluster's voxel are noted. Results are thresholded at p<0.001 for voxel and FWE-corrected (p<0.05) for cluster size. Functional labels which are visible on the figures are marked in bold.

Table S12

Difference between flanker and Simon-related activity exceeding time-on-task effect masked by their conjunction.

| Coordinates MNI x | y | z | Cluster size | (% of cluster) AAL Region | Cluster pFWE | Cluster pFDR | Cluster p(uncor) | Peak T | Cluster label |
|---|---|---|---|---|---|---|---|---|---|
| **(F0 > rtF0) > (S0 > rtS0) (mask with rtS0 ∩ rtF0)** | | | | | | | | | |
| 20 | 30 | 0 | 3362 | (81%) OUTSIDE<br>(6%) Caudate_R | < 0.001 | < 0.001 | < 0.001 | 6,96 | Caud<br>wm<br>**IPS** |
| 42 | 24 | -10 | 324 | (71%) Frontal_Inf_Orb_2_R<br>(14%) Insula_R<br>(6%) Frontal_Inf_Tri_R | < 0.001 | 0 | < 0.001 | 6,77 | **vlPFC**<br>+ aIns |
| -20 | -4 | 30 | 758 | (90%) OUTSIDE<br>(9%) Caudate_L | < 0.001 | < 0.001 | < 0.001 | 5,75 | Caud<br>wm |
| -32 | -70 | -28 | 760 | (51%) Cerebelum_Crus1_L<br>(32%) Cerebelum_Crus2_L<br>(7%) Cerebelum_6_L | < 0.001 | < 0.001 | < 0.001 | 5,2 | |
| -22 | -50 | 12 | 207 | (94%) OUTSIDE<br>(6%) Precuneus_L | 0 | 0 | < 0.001 | 5,16 | LOC L |
| 40 | -66 | -30 | 169 | (79%) Cerebelum_Crus1_R<br>(21%) Cerebelum_6_R | 0,01 | 0 | < 0.001 | 5,13 | |
| -32 | -68 | 12 | 372 | (77%) OUTSIDE<br>(11%) Occipital_Mid_L<br>(9%) Occipital_Inf_L | < 0.001 | < 0.001 | < 0.001 | 4,99 | wm<br>+ LOC L |
| -6 | -72 | -26 | 364 | (30%) Cerebelum_Crus2_L<br>(20%) Cerebelum_Crus1_L<br>(18%) OUTSIDE<br>(7%) Cerebelum_6_L<br>(6%) Vermis_8 | < 0.001 | < 0.001 | < 0.001 | 4,98 | |
| 38 | -90 | 0 | 350 | (34%) Occipital_Mid_R<br>(25%) Occipital_Inf_R<br>(24%) Calcarine_R<br>(13%) OUTSIDE | < 0.001 | < 0.001 | < 0.001 | 4,73 | **LOC R**<br>+ Cal R |

| | | | | | | | | | |
|---|---|---|---|---|---|---|---|---|---|
| 12 | 56 | 24 | 198 | (57%) Frontal_Sup_Medial_R<br>(42%) Frontal_Sup_2_R | 0 | 0 | < 0.001 | 4,28 | **dmPFC** |
| **(F0 > rtF0) < (S0 > rtS0) (mask with rtS0 ∩ rtF0)** | | | | | | | | | |
| 24 | -42 | -16 | 230 | (80%) Fusiform_R<br>(14%) Cerebelum_4_5_R | 0 | 0 | < 0.001 | 5,63 | **Fus R** |
| 50 | -60 | 4 | 210 | (97%) Temporal_Mid_R | 0 | 0 | < 0.001 | 5,4 | **pTmp R** |
| -34 | -50 | -22 | 335 | (82%) Fusiform_L<br>(12%) Cerebelum_6_L<br>(5%) Temporal_Inf_L | < 0.001 | < 0.001 | < 0.001 | 5,24 | **Fus L** |
| 28 | -60 | -16 | 191 | (81%) Fusiform_R<br>(13%) Cerebelum_6_R | 0 | 0 | < 0.001 | 5,12 | **Fus R** |
| -12 | -66 | 4 | 1087 | (28%) Calcarine_R<br>(25%) Lingual_R<br>(23%) Calcarine_L<br>(14%) Cuneus_L<br>(9%) Lingual_L | < 0.001 | < 0.001 | < 0.001 | 5,05 | **Cal**<br>+ Ling<br>**+ Cun L** |
| -8 | -58 | 64 | 174 | (80%) Precuneus_L<br>(19%) Parietal_Sup_L | 0,01 | 0 | < 0.001 | 4,78 | **Prec L** |
| -58 | -8 | 38 | 148 | (63%) Postcentral_L<br>(37%) Precentral_L | 0,01 | 0 | < 0.001 | 4,62 | **SM L** |
| -20 | -82 | 50 | 155 | (27%) Occipital_Mid_L<br>(25%) Parietal_Sup_L<br>(21%) Occipital_Sup_L<br>(13%) OUTSIDE<br>(12%) Precuneus_L | 0,01 | 0 | < 0.001 | 4,56 | **LOC L**<br>**+ Prec L** |
| 54 | -20 | 48 | 289 | (54%) Postcentral_R<br>(42%) SupraMarginal_R | < 0.001 | < 0.001 | < 0.001 | 4,47 | **SMG R**<br>**+ SM** |
| -56 | -24 | 16 | 133 | (47%) SupraMarginal_L<br>(32%) Temporal_Sup_L<br>(16%) Postcentral_L<br>(5%) Rolandic_Oper_L | 0,02 | 0 | 0 | 4,34 | **SMG L** |

Functional regions are reported with corresponding MNI coordinates of peak activity within each cluster as well as cluster size. Additionally noted is the percentage of cluster's voxel in each anatomical region defined using the AAL3 atlas. Only regions with more than 5% of all cluster's voxel are noted. Results are thresholded at $p<0.001$ for voxel and FWE-corrected ($p<0.05$) for cluster size. Functional labels which are visible on the figures are marked in bold.
