## supplementary figures for "Common and distinct BOLD correlates of Simon and flanker conflicts which can(not) be reduced to time-on-task effects"

### Supplementary materials include:

1) Supplementary figures S1-S5, included in this file

2) Supplementary tables S1-S12

3) Unthresholded and unmasked group-level whole-brain results maps are available at Neurovault repository: https://neurovault.org/collections/NTRDGLJW/.

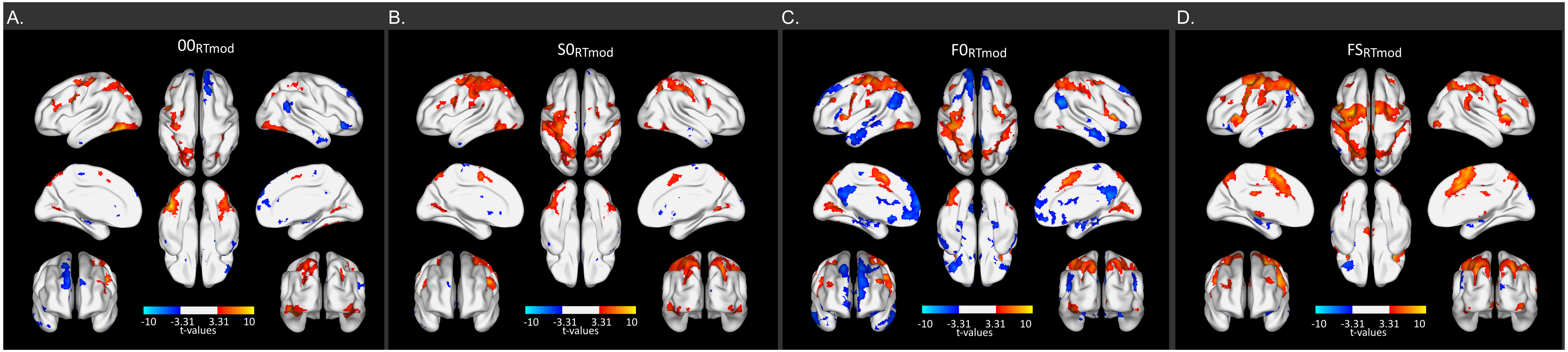


**Supplementary Figure S1. Activity correlated with RT variability within each of the task conditions.**

A) Map of the RT parametric modulation in no-conflict (00) condition. This map was used for calculating RT-predicted maps for each of the conflicts (rtS0, rtF0, rtFS). B-D) Maps of the RT parametric modulations in each of the conflict conditions.

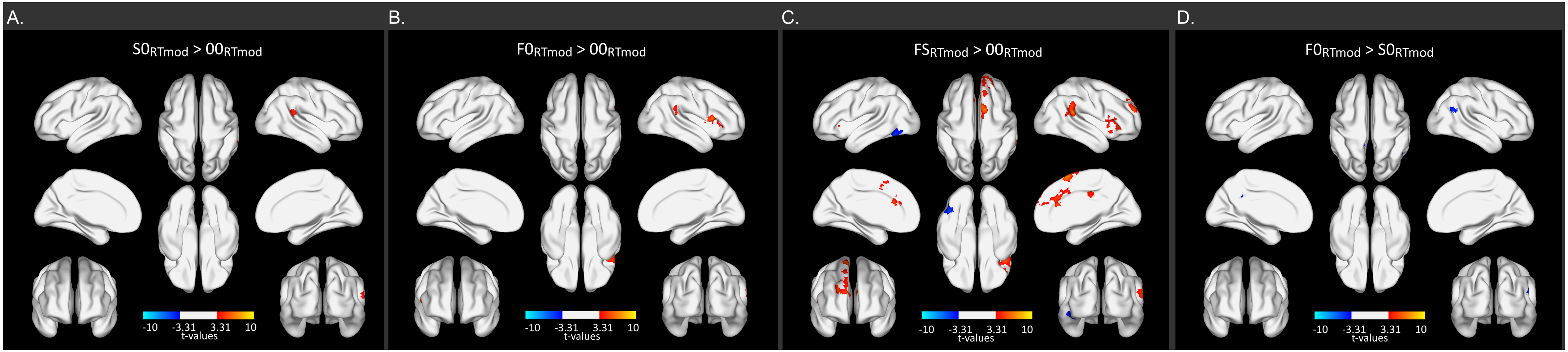


**Supplementary Figure S2. RT modulation difference between task conditions.**

Contrasts between: A) RT modulation within the Simon and no-conflict; B) flanker and no-conflict; C) multi-source and no-conflict D) flanker and Simon conditions. Contrast maps are thresholded at voxel-level $p<0.001$ and FWE-corrected ($p<0.05$) for cluster size.

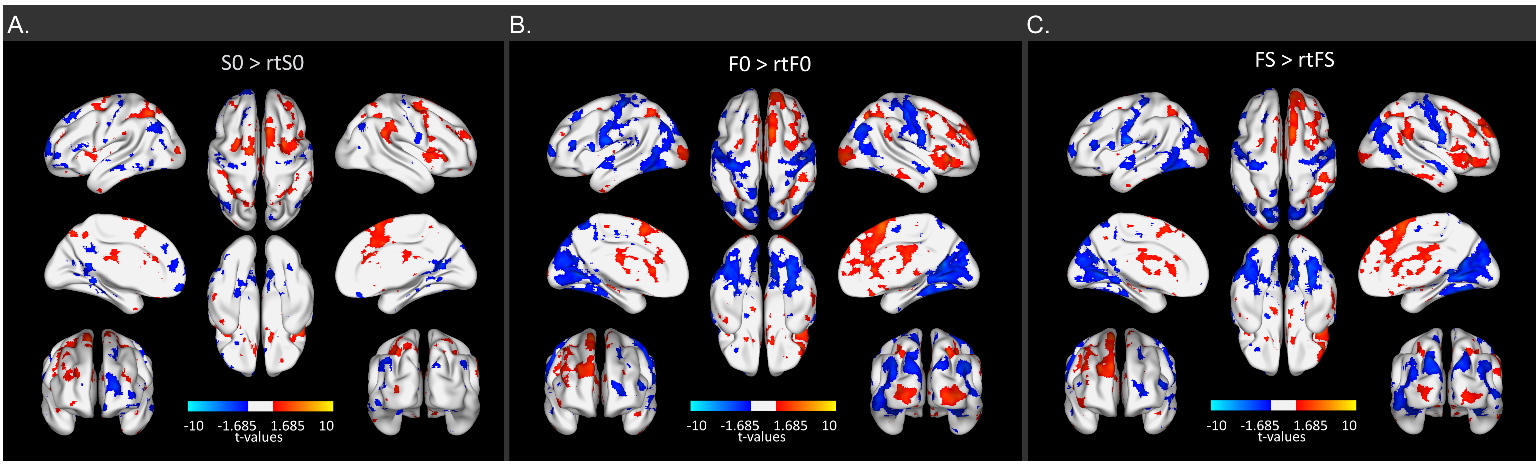


**Supplementary Figure S2. Regions with activations/deactivations exceeding the time-on-task effects** in the three types of conflicts induced by MSIT at a lowered significance level (voxel-level p<0.05), matching the ROI analysis threshold.
This figure is analogous to the main Figure 4.

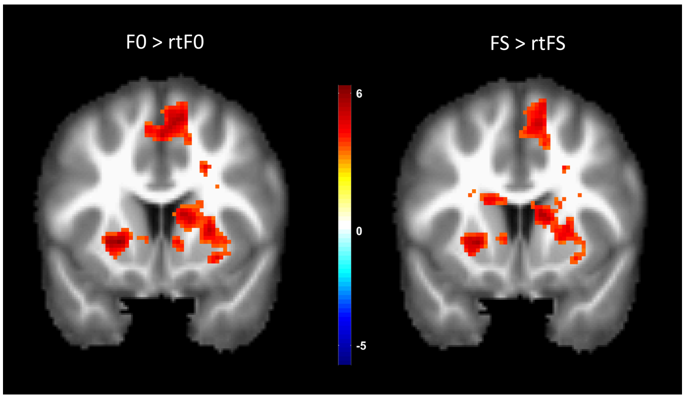


**Supplementary Figure S4. Subcortical activations exceeding the time-on-task effects.**
Volumetric view of the activation contrast map between observed and RT-predicted activity for F0 (left) and FS (right) conflict conditions. Coronal plane at y = 12. Note activity exceeding time-on-task in putamen, caudate and pMFC. There were no significant clusters of activity in S0 vs rtS0 comparison in subcortical areas.

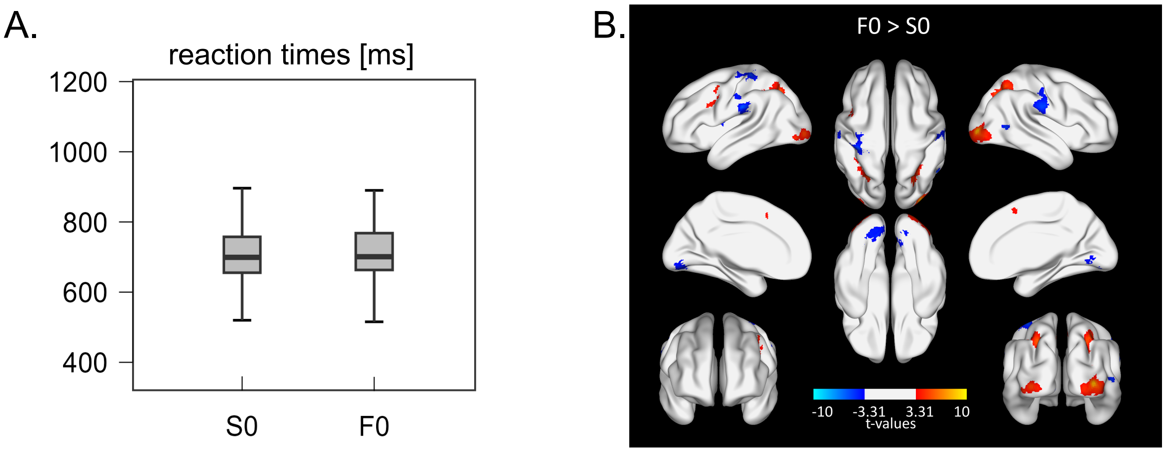


**Supplementary Figure S5. RT-matched analysis.** A) Reaction times in RT-matched S0 and F0 subsamples, y-axis set as in the main Figure 1B showing RT for the whole dataset. Note that the matching procedure required us to exclude ~23% slowest F0 and fastest S0 trials for each participant to nullify the statistical difference between F0 and S0 RT distributions (p = 0.48, Wilcoxon signed rank test). B) Difference between flanker- and Simon-related effects, when only RT-matched trials were selected from both categories. This figure can be compared with the main Figures 3B and 5B.
